# A guide RNA repeat checkpoint steers CRISPR-Cas9 catalysis

**DOI:** 10.64898/2025.12.02.691602

**Authors:** Ramadevi Chilamkurthy, Sruthi Sudhakar, Adrian A. Pater, Michael S. Bosmeny, Francis Stabile, Adam Katolik, S. Harikrishna, Rolf Turk, Masad J. Damha, Sabrina Leslie, Sergey Korolev, P.I. Pradeepkumar, Keith T. Gagnon

## Abstract

A widely adopted CRISPR-Cas9 modification is fusion of the naturally occurring two-component dual guide RNA (dgRNA) to create an artificial single guide RNA (sgRNA). However, mechanistic and functional differences between dgRNA and sgRNA have not been systematically explored. By investigating the activity of these two guide architectures, we discover a guide RNA repeat checkpoint (GRC) that senses the structure and dynamics of the guide repeat region. The GRC coordinates with other checkpoint mechanisms known to recognize spacer-target R-loop fidelity, together licensing target cleavage. Based on these principles, dgRNA and sgRNA properties could be combined into guide repeat-truncated sgRNAs (grtRNAs) and paired with high-fidelity Cas9 variants to further reduce off target editing. A mechanism that helps govern Cas9 catalysis via guide RNA structure sensing and communicates with other checkpoints offers a previously unappreciated path for understanding and further improving gene editing outcomes.

## INTRODUCTION

Cas9 from *Streptococcus pyogenes* (SpCas9) is a prototypical CRISPR-associated effector nuclease used for gene editing in biotechnology and therapeutics ^1–14^. SpCas9 evolved to naturally use two distinct RNAs, CRISPR RNA (crRNA) and *trans*-activating crRNA (tracrRNA), to guide DNA cleavage ^15–19^. An early modification was fusion of the dual guide RNA (dgRNA) system into an artificial single guide RNA (sgRNA) architecture ^20^, which simplified design and delivery. The use of sgRNA has become the conventional approach ^18,21–27^. Conversely, dgRNA has often been considered less efficient ^18,20,28^ despite successful use and commercialization ^29–33^. Consequently, differences in activity or mechanism between these two guide RNA architectures has not been systematically investigated ^18,20,25,34,35^.

Despite extensive characterization of SpCas9, no high-resolution structures or molecular dynamics studies with dgRNA have been previously reported and tools to predict the best SpCas9 guides for editing efficiency and specificity are trained with sgRNA data ^23,24,36,37^. However, some applications may benefit from dgRNAs^38^. For example, chemical modification of larger sgRNAs is more resource-intensive compared to separate crRNAs and tracrRNAs ^30,32,39^, which could enable certain CRISPR-based therapeutics ^29,40,41^. Studying SpCas9 guided by dgRNA may also uncover new mechanisms of regulation or provide insight into molecular evolution or CRISPR biology ^38^.

In this study, we performed a systematic comparison of SpCas9 activity when guided by dgRNA or sgRNA. We found that dgRNAs often performed as well or better across multiple spacer sequences and usually induced lower off target editing. By using model spacers and biochemical, structural, and computational methods, as well as SpCas9 mutations and guide RNA modification, we identified a previously uncharacterized mechanism, which we refer to as the guide RNA repeat checkpoint (GRC). The GRC senses SpCas9 interaction dynamics with the guide RNA repeat, the only structural difference between sgRNA and dgRNA. Our results support the coordination of the GRC with other checkpoint mechanisms involved in R-loop sensing, rationalizing the codependence of spacer sequence and guide architecture in steering SpCas9 catalysis. Combining both guide architectures into a single guide repeat-truncated sgRNA design, called grtRNA, created a smaller sgRNA that predictably reduced off target editing. Some high-fidelity variants of Cas9, which bear mutations designed to modulate the REC and bipartite seed checkpoint mechanisms, could be paired with grtRNAs to further enhance specificity. Thus, guide RNA interactions with Cas9 GRC residues represent a new mechanism that can be explored, in combination with other known checkpoint mechanisms, to create higher fidelity enzymes.

## RESULTS

### Guide architecture controls Cas9 activity

To initially compare editing between dgRNA and sgRNA (**Fig. 1A**), we electroporated Cas9 ribonucleoproteins (RNPs) assembled with fifteen dgRNAs or sgRNAs. Guides were selected based on a previous screen using commercial chemically modified guides ^42^ where five sgRNAs edited better, five dgRNAs edited better, and five performed equivalently (**fig. S1A**). When tested as unmodified guides, we observed six that performed better as dgRNAs, five with similar editing, and four that were better as sgRNAs (**Fig. 1B**). These results confirmed that dgRNAs can sometimes perform better than sgRNAs and underscored spacer-dependent differential activity between architectures (**fig. S1, B and C**).

**Figure 1.**
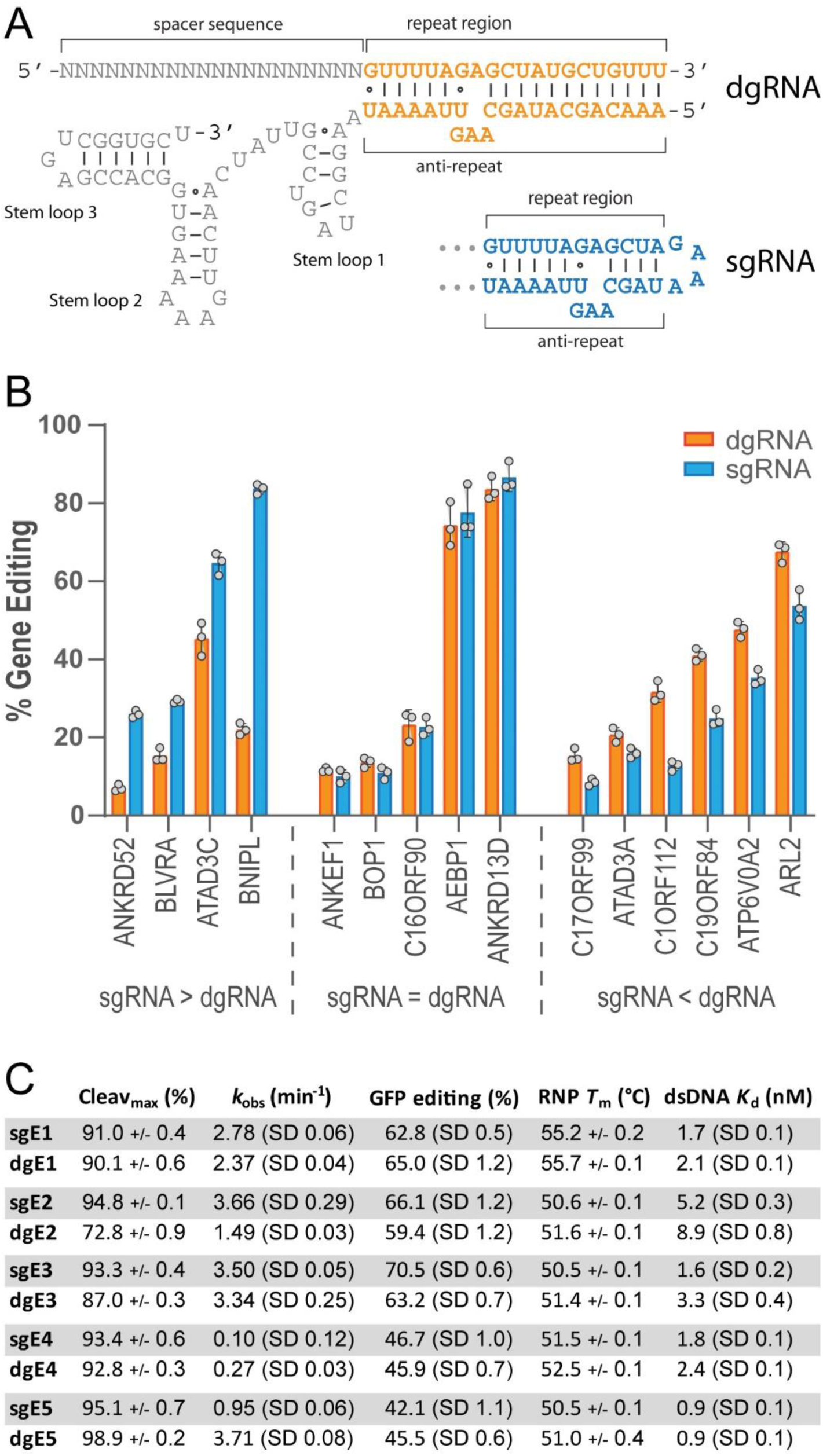
Single and dual guide Cas9 complexes exhibit differential activity depending on the guide spacer sequence. (**A**) Schematic representation of dual guide RNA (dgRNA) and single guide RNA (sgRNA) secondary structure. (**B**) Cell-based editing at 15 endogenous loci with matched single guide (sg) or dual guide (dg) CRISPR-Cas9. Error is reported as standard error of the mean (S.E.M.) of n=3 experimental replicates. (**C**) Summary of *in vitro* cleavage activity (Cleav_max_ and *k*_obs_) (n=2), editing of EGFP reporter gene in cells (n=4), ribonucleoprotein thermal stability (RNP *T*_m_) (n=2), and target DNA binding affinity (dsDNA *K*_d_) (n=2) for matched sg and dg CRISPR-Cas9 complexes using five model spacers, E1 to E5. Range of values (+/−) or standard deviation (SD) are reported.

To further explore differential activity, we selected five new spacers, E1 through E5, that target EGFP ^32,39,43^ (**Fig. 1C**). We performed *in vitro* cleavage assays, determining maximum cleavage at 60 min (Cleav_max_) and apparent observed rate constants (*k*_obs_), and gene editing via EGFP knockout (**Fig. 1C**). E1 and E4 guides displayed nearly identical cleavage kinetics and EGFP editing for both guide architectures, though E4 yielded slower kinetics (**Fig. 1C**, and **fig. S2**). E2 and E3 sgRNAs showed higher cleavage and EGFP editing (**Fig. 2, A** and **B,** and **fig. S2**) while dgRNA was more active for E5 (**fig. S2**). The E2 guide showed the greatest divergence in activity, highlighting the impact of guide RNA architecture on Cas9 function (**Fig. 2, A** and **B**).

**Figure 2.**
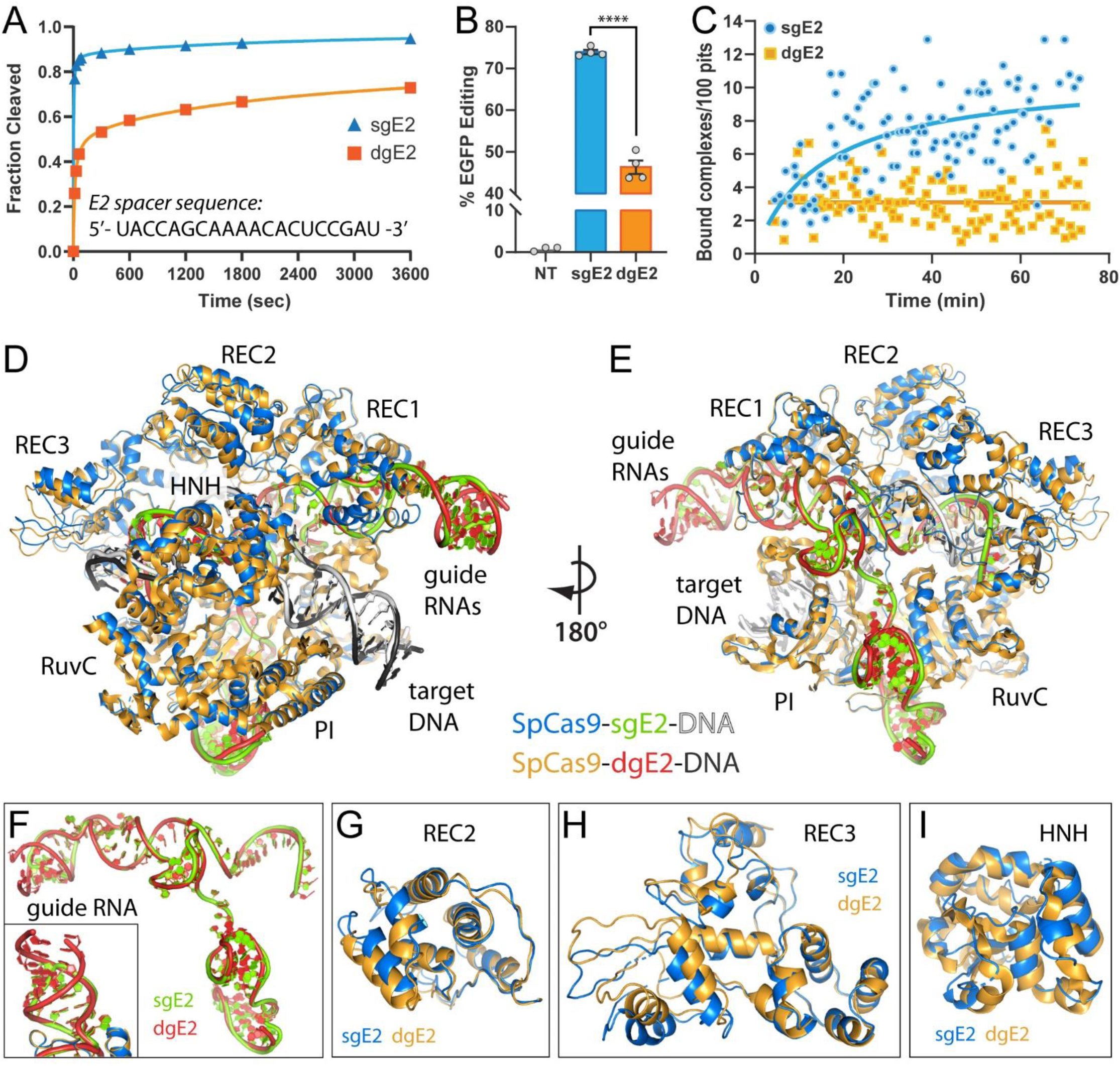
Activity, binding, and cryo-EM reconstructions of Cas9 ternary complexes guided by sgRNA and dgRNA with an E2 spacer. Comparison of sgRNA- and dgRNA-guided Cas9 with an E2 spacer sequence in (**A**) *in vitro* time course cleavage assays (n=2) and (**B**) cell-based EGFP editing assays (n=4). Error is reported as S.E.M. Unpaired t-test p-value: **** > 0.0001. (**C**) Target dsDNA binding measured by single-molecule convex lens induced confinement (CLiC) microscopy (n=2). (**D**) Ternary sgE2 (PDB ID: 8G1I) and dgE2 (PDB ID: 8FZT) complexes are shown superimposed and (**E**) rotated 180°. The single and dual guide RNAs are shown in (**F**) with a close-up of the GNRA tetraloop of sgE2 and the open stem of dgE2 shown in the inset. The REC2 (**G**), REC3 (**H**), and HNH (**I**) domains from the global superimposition are shown.

Single versus dual guides assemble two and three component RNPs, respectively, suggesting global complex stability may affect activity. Therefore, we compared RNP melting temperatures (*T*_m_) for Cas9 with E1 through E5 guides. However, RNP thermal stability was quite similar across guides (**Fig. 1C**, and **fig. S3, A** to **E**). We further performed limited trypsin hydrolysis of Cas9 bound to sgE2 or dgE2 and a complementary DNA target ^44^ but observed no major changes in cleavage patterns (**fig. S4**). These results suggest that global stability or large structural changes do not drive differential activity ^45^.

To test whether guide architecture alters target binding affinity, we determined dissociation constant (*K*_d_) values for target DNA (**Fig. 1C** and **fig. S3, F** to **J**). Though *K*_d_ values from bulk equilibrium binding also did not reveal clear differences, dgE2 binding affinity was notably reduced (**Fig. 1C**). Since the E2 spacer elicits the greatest differential Cas9 activity, we selected it for single-molecule binding studies. Using convex lens induced confinement (CLiC) ^46^, we found that sgE2 RNP binding increased over time while dgE2 RNPs achieved maximum yet incomplete binding within a few minutes, by the start of data collection (**Fig. 2C**). Thus, dgE2 RNP binding saturated quickly and incompletely while sgE2 RNP binding continued, suggesting that a fraction of dgE2 RNPs become trapped in non-productive conformations.

To determine if stability in the local guide repeat region affects activity, we chemically modified the tracrRNA of dgRNAs with two locked nucleic acid (LNA) residues near the 5’ terminus, thereby stabilizing crRNA-tracrRNA pairing (**fig. S5A**). The global stability for Cas9 RNP with modified tracRNA was unchanged (**fig. S5B**) yet conferred EGFP editing efficiencies that trended toward that of the sgRNA, typically falling between dgRNA and sgRNA (**fig. S5C**). These results suggest that differential activity is linked to repeat structure stability or dynamics.

### Guide architecture modulates conformational dynamics

To investigate more nuanced conformational changes across guide architectures, we solved matched cryo-EM structures of sgE2 and dgE2 Cas9 ternary complexes. A dgE2 complex was resolved to 3.03 Å while a sgE2 complex was resolved to 3.12 Å (**fig. S6, A** and **B**). Refined structures of each complex appear quite similar with a global root mean square deviation (RMSD) of 1.8 Å^2^ (**Fig. 2, D** and **E**) and both are similar to commonly observed “closed” (catalytically competent) conformations ^21^. The density corresponding to the repeat terminus of dgE2 was visible for five additional base pairs, as expected (**Fig. 2F**). Domains with visible shifts between sgE2 and dgE2 were REC2, REC3, and HNH (**Fig. 3, G to I**). These domains are known to act together through conformational checkpoints to sense proper R-loop structure, discriminate mismatches, and license catalysis ^47–51^, suggesting that guide architecture may impact specificity.

**Figure 3.**
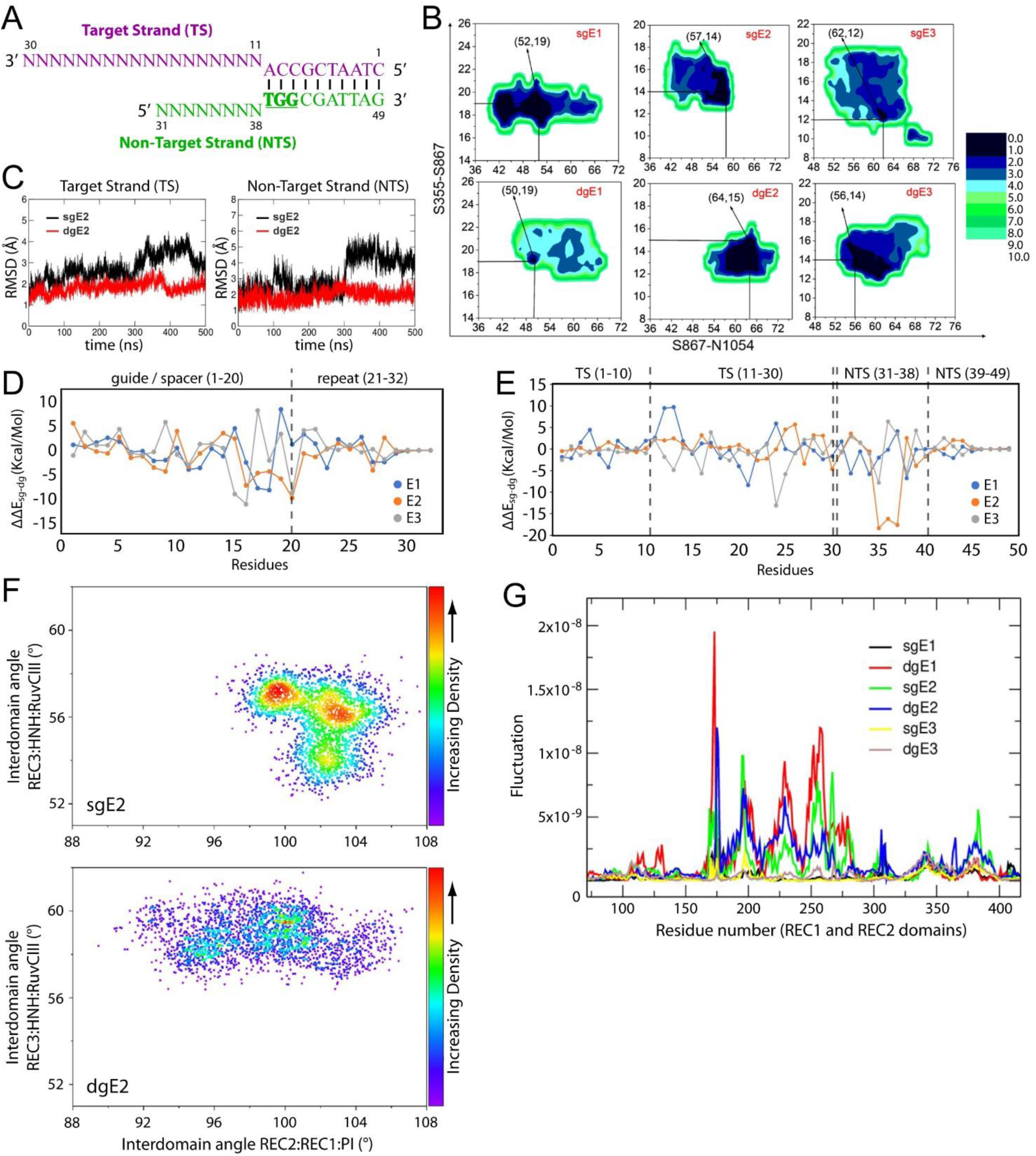
Molecular dynamics simulations identify energetics of target strand (TS) and non-target strand (NTS) DNA, dynamic REC domain motions, and fluctuations in REC domain residues that differ for sgRNA and dgRNA ternary Cas9 complexes. (**A**) Schematic illustration of the target strand (TS) and non-target strand (NTS) and their complementary base pairing used in MD simulation modeling. Numbering convention is continuous from the 5’ end of the TS (nts 1-30) to the 3’ end of the NTS (nts 31-49). (**B**) Potential mean of force (PMF) plots using S867-S355 and S867-N1054 amino acid distances in Cas9 calculated by reweighting 500 ns GAMD trajectories in AMBER 18. The coordinates corresponding to PMF = 0 are indicated by an arrow. (**C**) RMSD graphs of the DNA backbone of the TS (left) and NTS (right) within sgE2- and dgE2-guided Cas9 ternary complexes. RMSD values were calculated from the 500 ns GAMD trajectories. (**D**) Per residue energy decomposition plot of the repeat structure (crRNA:tracrRNA pairing region) of sgRNA and dgRNA in Cas9 ternary complexes. (**E**) Per residue energy decomposition plot of the TS and NTS DNA within the single and dual RNA guided Cas9 ternary complexes. For both panels **D** and **E**, ΔΔEsg-dg = ΔEsg − ΔEdg, ΔEsg is the per residue energy of the bases in the sgRNA and ΔEdg is the per residue energy of the bases in the dgRNA. Energy of Base X = sum of internal energy of base X + sum of interaction with bases + sum of interaction with amino acid residues (cut-off of 6 Å). (**F**) Density plots between the REC3:HNH:RuvCIII and REC2:REC1:PI domains of sgE2 and dgE2 ternary complexes. (**G**) Mobility plot of the REC1 and REC2 domains from visualizing principal component projections in VMD. All calculations were performed using the 500 ns GaMD trajectories.

Particles of the sgE2 complex selected for all high-resolution 2D classes yielded a single, well-refined 3D map (**fig. S6A**). However, data for the dgE2 complex yielded an additional prominent high contrast 2D class for nearly half of the particle images (**fig. S6C)**, which corresponded to an alternative class with the map refined to 4.2 Å. Resolution remained low at 3.87 Å despite attempts to improve resolution with additional data (**fig. S6D**). An available “open” conformation structure ^52^ can be partially fit to the alternative dgE2 complex density map (**fig. S6E**), suggesting that about half of the 2D classes observed may be trapped in conformations similar to open, catalytically incompetent states.

To gain insight into energetics and dynamics, we performed Gaussian accelerated molecular dynamics (GaMD) simulations with modeled E1, E2, and E3 sgRNA or dgRNA ternary Cas9 complexes ^53,54^ (**Fig. 3A**). We generated potential mean of force (PMF) plots, which depict low energy conformation sampling during simulations, using Cα distances between HNH and RuvC amino acids previously characterized by FRET ^55^ (**Fig. 3B**). The point of lowest energy (PMF = 0) occurred in the region of 50-65 Å (S867-N1054) and 12-20 Å (S867-S355) (**Fig. 3B**, and **fig. S7**) as expected ^56^, sampling of distances corresponded with the catalytic state of Cas9 (**fig. S7**), and Cas9 was relatively stable in all complexes based on root mean square deviation (RMSD) calculations (**fig. S8**). While RMSD values for guide RNA spacers and target strand (TS) and non-target strand (NTS) DNA fluctuated modestly for all complexes (**fig. S9** and **fig. S10**), E2 complexes showed significant differences in TS and NTS RMSD at 300 ns (**Fig. 3C**). We calculated root mean square fluctuation (RMSF) values for spacer RNA-TS heteroduplexes (R-loops), which were consistently low at ∼2 Å (**fig. S9D**), and the DNA target, which was much higher at up to ∼7 Å for the NTS of dgE2 (nts 31-33) (**fig. S11**). These results suggest that altered dynamics in the TS and NTS contribute to differential activity of model dgE2 and sgE2 complexes.

To better understand energetic differences, we performed molecular mechanics-generalized Born surface area (MM/GBSA) analysis. Fluctuations in energy differences for all guide RNAs appeared in the spacer seed region (nts 14-20) and beginning of the repeat structure (nts 21-32) (**Fig. 3D**). Significant energy fluctuations were also observed in the TS (nts 11-30) and the NTS (nts 31-38) (**Fig. 3E**). These fluctuations suggest a source of sequence-dependent energetic differences between guide architectures. Box plot representations of the geometric mean of hydrogen bond interactions between target DNA and Cas9 showed large differences between guide architectures for E2 (**fig. S12A**). Using the comparison circle algorithm, which takes into account H-bonds, interdomain angles, and RMSD of DNA strands to find overall variation, high variation was found for E2 complexes (**fig. S12B**).

To investigate Cas9 domain dynamics, we compared interdomain angles surrounding repeat structure (REC2:REC1:PI) (**fig. S13, A** and **B**) versus R-loop structure (REC3:HNH:RuvC-II) (**fig. S13, C** and **D**). The distribution of angles was more scattered for dgRNAs, indicating more dynamic domain movement (**Fig. 3F** and **fig. S13, E** and **F**). In particular, changes in the REC2:REC1:PI angles were more pronounced for dgRNA. These suggest that differences in guide architectures may be sensed during early target DNA binding by the PI and REC1 domains, which would be transmitted to other domains. Differences in interdomain angle values for E1, E2, and E3 also indicate spacer sequence- and repeat structure-dependent protein dynamics. Principal component analysis (PCA) of Cas9 using Cα atoms showed changes for both guide architectures of the same spacer, reinforcing structure-specific effects (**fig. S14**). The residue mobilities in Principal Component 1 (PC1) indicate that most fluctuations are experienced by the REC1 and REC2 domains (**Fig. 3G**), which confirms a conformational sensitivity of these domains to guide RNA architecture.

### Repeat structure and spacer sequence are codependent

To distinguish the relative contribution of TS or NTS sequence, we generated synthetic DNA targets where the NTS was varied to that of E1 through E5 or randomized, which should uncouple their contributions. Cas9 guided by dgE2 consistently cleaved all targets at 70-80% of sgE2 irrespective of the variable NTS sequence, reflecting the cleavage of a normal E2 target (**Fig. 4A** and **fig. S15A**). In contrast, using an E1 or E5 TS combined with the E2 NTS resulted in similar cleavage levels for both sgRNA and dgRNA, reflecting the typical cleavage of normal E1 and E5 targets (**Fig. 4B** and **fig. S15B**). These results suggest the TS, which interacts with the spacer to form R-loop structure, rather than the NTS plays a greater role in driving activity levels for each guide architecture.

**Figure 4.**
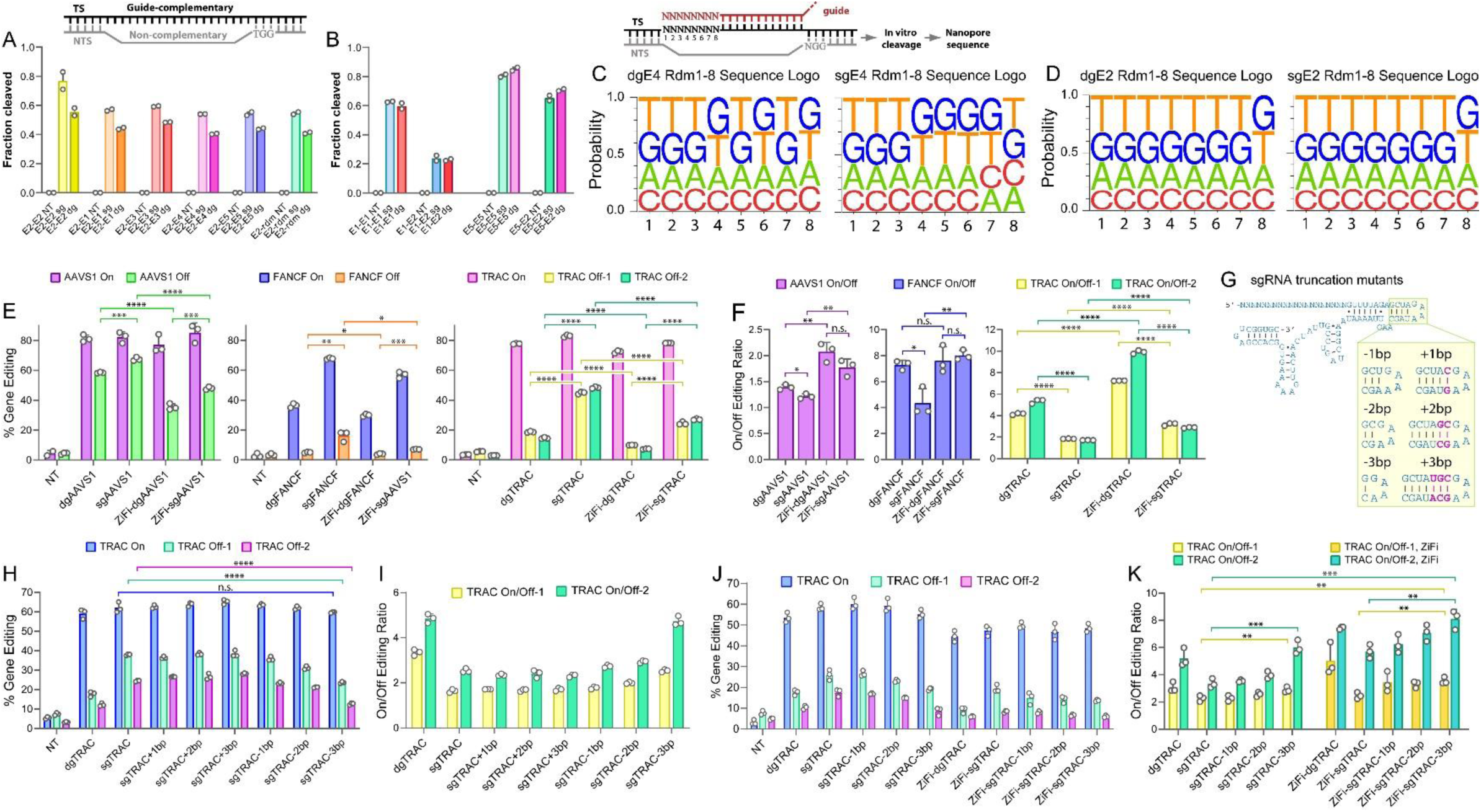
Target strand contribution, sequence preference, and specificity of Cas9 guided by sgRNA and dgRNA. *In vitro* cleavage assays where a TS complementary to the E2 spacer sequence (**A**) or E1 and E5 spacer sequences (**B**) were held constant while the NTS was varied. For example, E2-E1 indicates an dsDNA with an E2 TS and an E1 NTS. E2-rdm indicates a randomized NTS sequence. n=2 replicates. (**C-D**) Dual randomization of the 8 PAM-distal nts of guide RNA and target DNA to identify potential sequence preferences for Cas9 guided by dgRNA versus sgRNA. Products from *in vitro* cleavage were sequenced by nanopore to identify nt preferences at each randomized position. Average sequence logos from n=2 experimental replicates are shown across the 8 randomized positions for dgRNA and sgRNA. (**E**) Gene editing at AAVS1, FANCF, and TRAC genes and off target loci quantified by nanopore amplicon sequencing and CRISPResso (n=3). Dual guide and single guide RNAs were compared with and without ZiFi-Cas9. (**F**) On/off-target ratios determined from quantified gene editing data shown in panel E. (**G**) Illustration of the sequence-level mutations used to create stem mutants of sgTRAC. (**H-K**) Gene editing and on/off ratios at the TRAC gene and two off-target sites with sgTRAC stem mutants quantified by nanopore amplicon sequencing (n=3). Error is reported as S.E.M. Unpaired t-test p-value: * > 0.05, ** > 0.01, *** > 0.005, **** > 0.0001.

To explore whether guide architecture induces a PAM distal sequence preference for Cas9, we performed cleavage assays and deep sequencing of cleavage products using randomized PAM distal regions for E4 and E2 guides with matched, randomized targets (**Fig. 4, C** and **D**, and **fig. S15, C** and **D**). Full randomization of the entire 20 nt spacer abrogated detectable activity (**fig. S15C**). Nanopore sequencing of cleavage products resulted in sequence logos that showed only slight variations in nucleotide preferences between guide architectures (**Fig. 4, C** and **D**, and **fig. S16, A** to **D**), albeit with some bias observed at position 8. Together, these results suggest that spacer sequence influences guide architecture activity through remaining PAM proximal nts or a cumulative effect of the entire spacer sequence ^35,57–61^.

### Repeat structure influences specificity

To determine if Cas9 specificity is altered between guide architectures, we targeted three endogenous genes, AAVS1, FANCF, and TRAC-tgt15 (TRAC), which includes four previously characterized off target sites in the genome ^50^. dgRNAs offered similar on target editing as sgRNAs at AAVS1 and TRAC (**Fig. 4E**), as well as higher specificity profiles (on/off target ratio) at all off-target sites (**Fig. 4F**). To explore a possible connection with allosteric regulatory mechanisms, we combined two key mutations from the LZ3-Cas9 high-fidelity enzyme ^62^, T769I in the RuvC-HNH linker 1 and G915M in HNH-RuvC linker 2, with the REC3 domain mutation R691A found in HiFi-Cas9 ^63^. Average on-target editing with this new mutant, which we refer to as ZiFi-Cas9, was slightly reduced across all guides (**Fig. 4E**), including E1, E2, and E3 guides (**fig. S17, A** and **B**), but provided reductions in off target editing (**Fig. 4E**). Notably, improvements in on/off target editing by dgRNA were generally additive with ZiFi-Cas9 (**Fig. 4F**).

To investigate whether increased flexibility or dynamics of dgRNA was responsible for lower off target editing, we truncated the stem adjacent to the GAAA tetraloop in sgTRAC to reduce stability (**Fig. 4G**). While adding base pairs had no effect, truncating the stem lowered off target editing while largely preserving on target editing, mimicking dgTRAC (**Fig. 4H**). On/off target editing ratios for sgTRAC-3bp improved by about 1.3-fold and 1.85-fold with Cas9 and about 1.55-fold and 2.4-fold when paired with ZiFi-Cas9 at off-target sites 1 and 2, respectively (**Fig. 4, I** to **K**). For TRAC gene editing, the −3bp truncation showed no loss of on target editing (**Fig. 4J**).

### A guide repeat checkpoint helps regulate Cas9

Specificity modulation by dgRNAs or guide repeat-truncated sgRNAs, referred to as grtRNAs from here forward, may employ a mechanism that is distinct from, but overlapping with, R-loop structure-sensing mechanisms used by high-fidelity variants like ZiFi-Cas9. The guide repeat structure appears to act as a nexus for stabilizing interaction between Cas9 REC1 and PI domains, clasping around the RNA duplex and potentially serving as a type of RNP assembly checkpoint (**Fig. 5, A** and **B**). Perturbation in the stability or dynamics of this clasp structure might influence catalysis, which is supported by dgRNA and grtRNA activity. Therefore, we compared low-energy conformers from GaMD simulations for sgE2 and dgE2 complexes. We observed a π-π interaction between Y1131 and F105, a slightly parallel offset π-π interaction between W1126 and F105, and an H-bond between Y1131 and V107 in the sgE2 complex (**Fig. 5C**). However, in the dgE2 complex the partial π-π interaction between F105 and W1126 is absent and the H-bond between Y1131 and V107 is lost (**Fig. 5D**). Thus, the reduced catalytic activity of dgE2 might be associated with reduced stability of the GRC clasp.

**Figure 5.**
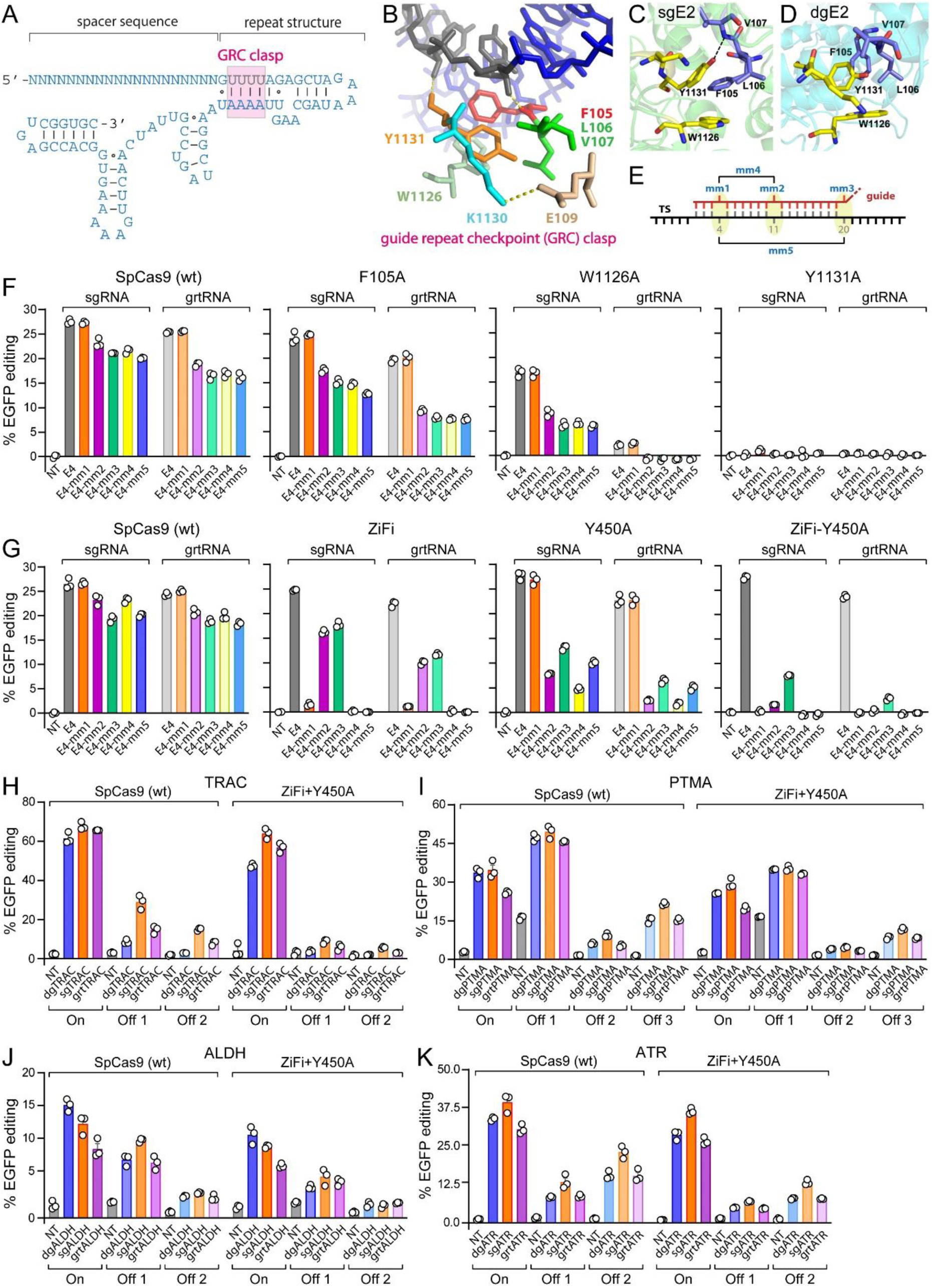
GRC mutants and high-fidelity Cas9 variants combined with grtRNAs modulate on and off target editing. (**A-B**) Illustration of GRC interaction with the guide RNA repeat structure. (**C-D**) GRC amino acid interactions observed in GaMD simulations for for sgE2 and dgE2 complexes. (**E**) Illustration of mismatched guide pairing to targets for off target assessment. (**F**) Editing of EGFP with the E4 spacer and mismatch-containing guides and Cas9 GRC mutants measured by flow cytometry (n=3). (**G**) Editing of EGFP with the E4 spacer and mismatch-containing guides and high-fidelity ZiFi, Y450A, and ZiFi-Y450A Cas9 variants measured by flow cytometry (n=3). (**H-K**) Editing of endogenous loci with guides targeting the TRAC, PTMA, ALDH, and ATR genes and the high-fidelity ZiFi-Y450A-Cas9 variant. On and off target site editing was measured by nanopore amplicon sequencing and CRISPResso2 (n=3).

To confirm the importance of the guide repeat clasp, we created three individual Cas9 mutants, F105A, W1126A, and Y1131A, which should disrupt hydrophobic packing (**Fig. 5, B** to **D**). These were paired with single mismatch-containing sgRNAs and grtRNAs, which mimic off target editing at five different positions along the spacer (**Fig. 5E**). F105A slightly reduced on target editing while offering small but significant reductions in off target editing (**Fig. 5F**). Combining F105A with grtRNA slightly increased on/off target ratios, though on target editing was reduced. W1126A showed a significant decrease in overall activity, losing nearly all editing when paired with grtRNA, while Y1131A completely abrogated editing (**Fig. 5F**). These results demonstrate that the guide repeat clasp serves as a checkpoint for licensing Cas9 catalysis.

Synergism between ZiFi-Cas9 and grtRNA suggested that multiple checkpoint mechanisms might be combined to potentially boost Cas9 fidelity. ZiFi mutations are designed to engage the REC domain checkpoint, which primarily senses PAM-distal mismatches ^47,49,50,63^. In support of this hypothesis, ZiFi-Cas9 nearly completely suppressed editing by mm1 (PAM-distal) sgRNA, yet only moderately reduced mm2 (PAM-central) and mm3 (PAM-proximal) sgRNA editing (**Fig. 5G**). In contrast, the bipartite seed checkpoint is proposed to sense PAM-central or PAM-proximal mismatches via direct spacer interactions with tyrosine 450 (Y450) ^64,65^. Indeed, a Y450A mutant had no effect on mm1 sgRNA editing but substantially reduced mm2 and mm3 sgRNA editing (**Fig. 5G**). When ZiFi-Cas9 and Y450A were individually paired with grtRNA, mm2 and mm3 editing was further reduced, but not mm1 editing (**Fig. 5G**). These results suggest that grtRNA, operating through the guide repeat checkpoint, referred to as the GRC from here forward, may primarily participate in discriminating PAM-proximal mismatched targets.

Strikingly, when a single variant combining ZiFi and Y450A mutations was paired with grtRNA, editing by mm1, mm2, and mm3 was almost entirely suppressed while on target editing was maintained at about 90% of normal Cas9 and sgRNA (**Fig. 5G**). Combining Y450A with individual or double ZiFi mutations did not generate better results (**fig. S18, A** and **B**). Editing by HF1, a previously characterized high-fidelity variant, showed similar off target editing as ZiFi-Y450A but greater loss of on target editing (**fig. S18C**). Testing with different spacers against EGFP, the E1 and E2 sequences, also showed reduced off-target editing with ZiFi, Y450A, ZiFi-Y450A, and HF1 (**fig. S19**). However, results varied across spacers and grtRNAs improved PAM-proximal off-target editing with mm3 with less pronounced effects than with the E4 guide, underscoring the variable spacer sequence dependence of checkpoint mechanisms in general.

To assess on and off target editing at endogenous loci, we paired dgRNA, sgRNA, and grtRNA with ZiFi-Y450A-Cas9 and targeted the TRAC, ALDH1A3 (ALDH), ATR, and PTMA genes ^62^ (**Fig. 5, H** to **K)**). On target editing was similar across guide architectures, though dgRNA performed best for the ALDH sequence and sgRNA best for ATR and TRAC. grtRNA generally reduced on target editing, except for the TRAC gene, but reduced editing at off target sites to a similar extent as dgRNA. dgRNA or grtRNA, paired with either Cas9 or ZiFi-Y450A-Cas9, reliably generated the best on/off target ratios. Together, these results support a role for the GRC in controlling Cas9 activity and demonstrate the potential to combine multiple checkpoint mechanisms for higher fidelity Cas9 editing.

## DISCUSSION

Although Cas9 is well-characterized among CRISPR-Cas enzymes, its regulation and mechanisms when guided by a natural dgRNA have been largely unexplored. Here, while comparing the activity of Cas9 guided by dgRNA and sgRNA, we noted that dgRNA often worked as well or better than sgRNA but performance depended on spacer sequence. Differential activity observed between the two guide architectures arose from the guide RNA repeat region and was traced to changes in target DNA and Cas9 structural dynamics, suggesting guide architecture could also influence specificity. Indeed, dgRNA and grtRNA were found to provide reductions in off target editing and these were additive with Cas9 variants bearing mutations designed to utilize the REC domain and bipartite seed checkpoints. These results support the identification of a new checkpoint mechanism, the GRC, which is comprised of a motif that joins the PI and REC1 domains around the guide repeat structure. The GRC has the potential to be tuned, either through mutagenesis of Cas9 or guide RNA modification, and combined with mutations of other checkpoint mechanisms to further enhance Cas9 fidelity.

Assembly of Cas9 RNP complexes involves docking of guide RNA onto the Cas9 protein ^20,25^. Recognition of the correct guide RNA creates an interaction nexus at the repeat structure, enabling the PI and REC1 domains to physically clasp around the RNA duplex. The GRC clasp motif is stabilized by a salt bridge, hydrogen bonding, and hydrophobic packing that is not resolved and therefore not observed in Cas9 apo structures ^52^. This clasp interaction is essential for activity, as demonstrated by GRC mutants like Y1131A, and likely serves as a key RNP assembly checkpoint. However, sensing of repeat structure and dynamics also influences catalysis. This effect is best explained by communication with other catalytic checkpoint mechanisms through allostery. Though the GRC clasp is quite distant from the HNH and RuvC catalytic centers, the PI and REC domains are both linked to target DNA engagement through PAM recognition and R-loop interaction, respectively, and are implicated in known allosteric activation mechanisms ^25,49,50,56,66–69^. Conformational shifts in the REC2, REC3, and HNH domains were apparent in matched ternary cryo-EM structures, supporting an allosteric link to catalytic efficiency and specificity.

The codependence of spacer sequence and repeat structure on Cas9 catalysis also suggests collaboration of the GRC with other checkpoints to ultimately license catalysis. MD simulations identified dynamics arising from the spacer, especially the seed region, the DNA TS and NTS, and Cas9 REC1 and REC2 domains centered around the PAM-proximal R-loop and repeat structure. The REC domain checkpoint senses R-loop integrity while the bipartite seed checkpoint, which utilizes Y450 in the REC1 domain, surveys PAM-proximal R-loop formation ^47,49,64^. Both checkpoint mechanisms are known to influence Cas9 specificity and can be modulated with amino acid mutations to increase catalytic fidelity ^62,64,65,70^. grtRNAs can provide additive improvements in specificity when paired with mutations that act through the REC domain checkpoint (ZiFi-Cas9) and bipartite seed checkpoint (Y450A-Cas9), suggesting distinct but interdependent mechanisms.

Altered dynamics can change the energetic landscape of Cas9 conformational transitions required to engage and cleave target DNA ^71,72^, which can induce lower off target editing by partially inhibiting allostery or catalysis ^69,73^. Many high-fidelity Cas9 variants function by mildly reducing target affinity or altering allosteric networks. For example, the HF1 variant of Cas9 was rationally designed to reduce the dwell time of Cas9 binding on target substrates by reducing charge-charge interactions with the R-loop structure ^65^. Mutations of variants selected by directed evolution are predicted to function by similar mechanisms ^70^. Unfortunately, these mutations often come at the cost of reduced on target editing efficiency. Combining multiple independent mechanisms might minimize off target editing and maximize on target editing ^62,69^.

A more flexible or less stable repeat region, such as with dgRNA or grtRNA, can increase or decrease cleavage and editing efficiency depending on the spacer sequence. These results would suggest an intrinsic sequence preference for dgRNA versus sgRNA, which would enable better predictability for guide architecture selection. However, partial randomization of the PAM-distal spacer-target region and sequencing of cleavage products for two separate spacer sequences did not reveal a clear sequence logo difference between sgRNA and dgRNA, suggesting preferences may lie in the PAM-proximal spacer region. Swapping non-complementary NTS sequences, combined with MD studies, also support a role for the PAM-proximal R-loop in coordinating with the GRC to dictate Cas9 activity and specificity. Nonetheless, we found that dgRNA and grtRNA usually conferred lower off target editing regardless of spacer sequence.

A commonly held assertion is that dgRNAs do not perform as well as sgRNAs. However, this study establishes that dgRNAs can sometimes be superior, making a compelling case to include dgRNAs in more studies and further investigate their specificity. Likewise, the creation of grtRNA helped test GRC mechanism and guide repeat dynamics while displaying decreases in off target editing. However, only a limited number of spacer sequences were investigated. We found that one spacer sequence, E2, showed substantial differential activity between sgRNA and dgRNA, nominating it as a model for detailed analyses. While E2 provided an informative example, results may not translate broadly to other spacer sequences. To fully explore the potential for the GRC to improve fidelity, deep mutagenesis or directed evolution techniques may be required for both the guide RNA repeat region and Cas9 protein. Nonetheless, combinations of GRC mutants and guide repeat modifications should enable even higher fidelity Cas9 variants when combined with other mutations, such as ZiFi-Y450A-Cas9 ^49,51,53,62,65,66^.

Roles for the guide RNA repeat structure and the GRC in Cas9 fidelity appear to be previously unknown or unappreciated. Amino acids directly involved in formation of the GRC clasp have not appeared in other high-fidelity Cas9 variants ^62,70^ and the guide RNA repeat sequence and structure have remained unchanged since the earliest published CRISPR-Cas9 studies ^20,25^. The grtRNA design introduced here is distinct from previously reported truncated guide RNAs ^74,75^, where the 5’ end of the spacer is simply shortened, as these modulate R-loop structure/stability sensing, presumably through mechanisms like the REC domain checkpoint.

The principles uncovered here provide a framework for developing new, more specific CRISPR-Cas tools and safer therapeutics. These include focused mechanistic studies on the effects of repeat structure on checkpoint mechanisms, optimization of guide repeat structures, and combining grtRNAs and GRC mutants with other high-fidelity strategies, like Cas9 variants ^62,70^ or truncated spacers ^74^. Our findings may also translate to other Cas9-based platforms, like base editing or prime editing where binding specificity is paramount or sgRNAs are necessary ^8,12,13^. Chemical modifications in the guide repeat region are expected to alter dynamics and their impact on Cas9 fidelity, especially for therapeutic applications, should be carefully considered ^41^. The grtRNA design also offers a more compact sgRNA and the potential for further optimization. Finally, the phenomenon of an RNP assembly checkpoint, where binding to a unique RNA structure or sequence is prerequisite for catalysis, suggests the potential for mechanisms like the GRC to impact efficiency and specificity broadly across other RNA-guided and CRISPR-Cas systems.

## METHODS

### RNA Synthesis

DNA oligonucleotides, T7 DNA templates for *in vitro* transcription, crRNAs, tracrRNAs and sgRNAs were chemically synthesized by Integrated DNA Technologies (IDT). Chemically modified tracrRNA was custom synthesized following our previously published protocols ^76,77^. Select tracrRNAs and sgRNAs were also prepared by T7 *in vitro* transcription with DNA templates following standard protocols ^78^. All RNA and DNA oligonucleotide sequences can be found in **Table S1**. Briefly, single-stranded DNA templates were annealed to a T7 promoter oligo by heating at 95°C for 3-5 min followed by slow cooling to generate double-stranded promoter regions, which support *in vitro* transcription by T7 RNA polymerase. Transcription reactions containing purified T7 RNA polymerase, 30 mM Tris (at pH 7.9), 40 mM MgCl_2_, 2% PEG 8000, 0.005% Triton X-100, 2 mM spermidine, 10mM DTT, SUPERase-in (Invitrogen), 5 mM NTP mix and 2.5 μM T7-DNA template were prepared and incubated at 37°C for 2 h. Afterward, the DNA template was degraded by the addition of 1 U of DNase I per 20 μL of reaction and incubated at 37°C for 15 min. Reactions were stopped by addition of 50 mM EDTA and phenol-chloroform extraction. The transcription products were separated by denaturing urea-PAGE and visualized using methylene blue staining. Full-length RNA products were extracted by gel purification using crush and soak elution and ethanol precipitation, quantified by absorbance at 260 nm and Beer’s law using calculated extinction coefficients (nearest neighbor approximations).

### Preparation of Cas9 Enzymes

Plasmid encoding *Streptococcus pyogenes* Cas9 (SpCas9) with a C-terminal fusion of a nuclear localization signal (NLS) and a 6x-Histidine tag (pET-Cas9-NLS-6xHis) ^79^ was obtained from Addgene (62933). A catalytically inactive “dead” Cas9 (dCas9) was prepared by performing site-directed mutagenesis on this plasmid to generate H840A and D10A mutations (pET-dCas9-NLS-6xHis). Cas9 proteins were prepared similarly as previously described ^80^ with some alterations. Briefly, protein expression was induced in Rosetta (DE3) cells with 0.4 mM IPTG at 18°C for 16 h. Cell pellets were resuspended in chilled binding buffer (20 mM Tris-HCl, pH 7.5, 250 mM NaCl, 1 mM PMSF, 5 mM imidazole). Resuspended cells were lysed by either sonication or French press and the resulting lysate was clarified by centrifugation. His-Pur Cobalt-CMA resin (Thermo Scientific) was equilibrated with binding buffer and the supernatant added and rotated at 4°C for 1 h. The supernatant was washed sequentially with at least three bed volumes each of increasing concentrations of NaCl in wash buffer (Tris-HCl, pH 7.5, 0.25/0.5/0.75/1.0 M NaCl, 10 mM imidazole). Protein was eluted with elution buffer (Tris-HCl, pH 7.5, 250mM NaCl, 130 mM imidazole). Purified Cas9 enzyme was concentrated and buffer-exchanged into gel filtration-running buffer (20 mM Tris-HCl, pH 7.5, 200 mM KCl, 0.5mM DTT) using Vivaspin 15 centrifugal concentrators (Sartorius, 30K MWCO). Concentrated protein was injected into pre-equilibrated Superdex 200 column and 1ml fractions collected. Fractions with Cas9 were concentrated, buffer exchanged in to 2x storage buffer (40 mM Tris-HCl, pH 7.5, 400 mM KCl, 1mM DTT), and an equal volume of glycerol added to obtain final of 50% glycerol in 1x storage buffer. Concentration was determined by UV absorbance at 280 nm using a calculated extinction coefficient (120,450 M-1 cm-1) ^81^ and Beer’s law.

### *In Vitro* Cas9 Cleavage Activity Assays

*In vitro* cleavage assays were performed as previously described ^32,43^. Two versions of EGFP target gene (EGFP, non-codon optimized; EGIP, codon optimized) were used to access the intrinsic enzyme activity of Cas9 loaded with different guide RNAs. Plasmid DNA containing the EGFP target gene (pSMART-EGFP-Cut1c) was linearized with either HindIII or KasI restriction enzymes and then purified by phenol-chloroform extraction and ethanol precipitation. A 1 kb fragment of the target EGIP gene was PCR-amplified using “pEGIP_vitro target” primers from pEGIP plasmid DNA (Addgene, 26777) and purified by phenol-chloroform extraction and ethanol precipitation.

Briefly, a master mix containing final concentrations of 0.75 µM Cas9, 0.25 µM tracrRNA, 0.3 µM crRNA or 0.75 µM Cas9 and 0. 25 µM sgRNA in a 1x cleavage buffer (20 mM Tris-HCl, pH 7.5, 100 mM KCl, 5% glycerol, 1 mM DTT, 0.5 mM EDTA, 2 mM MgCl_2_) supplemented with 0.1 mg/mL of purified yeast tRNA was incubated at 37°C for 10 min to allow the RNP to assemble. The concentration of tracrRNA or sgRNA was purposely set as the limiting component of the RNP complex and used to predict final RNP concentration. To each reaction tube containing 1µl of target DNA (100 ng/µL), 39µL of pre-assembled RNP was added at 37°C and reactions stopped at specified time points by addition of 2% LiClO_4_ in acetone and placed on ice. The samples were allowed to precipitate at −20°C for at least 1 h, pelleted by centrifugation and washed with acetone. Air dried pellets were resuspended in TE buffer or ddH_2_O and then treated with 10 μg of RNase A (Thermo Scientific) for 15 min followed by 20 μg of Proteinase K (Thermo Scientific) for 15 min at room temperature. Samples were mixed with 6X Purple Gel Loading Dye (New England Biolabs) and the cleavage products were resolved on 1% or 1.5% Tris-Borate-EDTA (TBE) agarose gels. Agarose gels were stained with ethidium bromide and visualized using a UV imager. The fractions of target cleaved were quantified using ImageJ software (v1.43u). The band intensity for the cleavage product band was divided by the combined intensity of cleavage product and uncut substrate bands and reported as fraction cleaved (i.e. “cut” / “cut + uncut”). Time course cleavage assay results were plotted using Prism (Graphpad) software fit to a one-site binding hyperbola equation, which was used to calculate maximum cleavage (Cleav_max_) and time required to achieve 50% Cleav_max_ (t_½_). Observed rate constants (*k*_obs_) were approximated by the equation *k*_obs_ = ln(2/t_½_) ^82^.

### Dot-Blot Filter Binding Assays for dCas9 Target DNA Binding

Synthetic crRNA and target double-stranded DNA lack 5’ phosphates and were directly radiolabeled. 100 pmols of tracrRNA, sgRNA, or target strand DNA was radiolabeled with [γ-^32^P]-ATP using T4 Polynucleotide Kinase (New England Biolabs) following the manufacturer’s recommended protocol. Reactions were phenol-chloroform extracted and radiolabeled RNA or DNA was gel-purified on 15% denaturing polyacrylamide gels (1x TBE, 7 M urea) by the crush-and-soak method. Gel-purified radiolabeled RNA and DNA was quantified by scintillation counting.

The active concentration of dCas9 was empirically determined as previously described ^83^. Briefly, increasing amounts of dCas9 were titrated with 40 pmol (1 µM final) of cold (unlabeled) crRNA:tracrRNA complex in which a negligible amount of radiolabeled crRNA (1,000 cpm) was spiked. Dot blots were performed to identify the volume of dCas9 corresponding to 40 pmol (complete binding of crRNA) in the reactions described. For target DNA binding by dCas9 RNP complexes, radiolabeled duplex target DNA (1,000-1,500 cpm/reaction) was combined with increasing concentrations of a pre-assembled dCas9-tracrRNA-crRNA or dCas9-sgRNA complex in a final reaction of 60 µL in 1x cleavage buffer (see above) and 0.1 mg/mL tRNA. After incubation at 37°C for 15 min, reactions were vacuum filtered over nitrocellulose membrane (Protran Premium NC, Amersham) using a 96-well dot blot apparatus. Wells were washed twice with 200 µL of 1x cleavage buffer. Membrane was then removed and washed with 1X phosphate-buffered saline (PBS) and air dried at room temperature. Binding of radioactive target DNA was then visualized by phosphorimager. Spots were quantified with ImageQuant software, plotted in Prism (GraphPad) and fit to a one-site binding hyperbola equation. Error bars for all quantified data represent experimental replicates, not technical replicates. Samples size was selected based on the expectation that three or more replicates will be representative.

### Single Molecule Target DNA Binding by dCas9

For RNP assembly, 36 nM of dCas9 and 30 nM of Cy3-sgE2 were mixed in 20 mM Tris-HCl (pH 7.5), 100 mM NaCl, 0.5 mM MgCl_2_, and 0.5 mM DTT. The sample was then incubated at 37°C for 10 min. For dual guide experiments, the Cy3-sgE2 was replaced with 30 nM of Cy3-crE2 and 30 nM of tracrRNA. While the RNP sample incubated, 21.1 nM of pSMART-EGFP-Cut1c plasmid ^43^ was mixed in 12 mM Tris-acetate (pH 7.5), 2.5 μM yeast tRNA, 0.2% Tween-20, 2.5 mM protocatechuic acid (PCA), 50 nM protocatechuate 3,4-dioxygenase (PCD), and 2 mM Trolox. The PCA/PCD oxygen scavenging system and Trolox are added for anti-bleaching and photostability. For control experiments, the pIDT-EGFP-Cut1c plasmid was replaced by an unrelated plasmid of similar molecular weight (pTRE3G plasmid), which does not include the EGFP target sequence.

Microscopy setup, data acquisition, and image analysis were performed using convex lens-induced confinement (CLiC) microscopy as described previously ^84^. After incubation, the RNP sample was diluted to 750 pM in the plasmid sample, mixed, and then flowed using positive air pressure into the CLiC device. The sample enters a flow cell consisting of two coverslips separated by 10 µm double sided spacer tape (Nitto Denko). The bottom coverslip contains an array of 3 µm diameter wells with 500 nm depth that were etched into the coverslip using microfabrication methods ^84^ while the top coverslip is featureless. In this microscopy technique, a convex lens pushes down on the top coverslip, deforming it, and bringing it into contact with the bottom coverslip. When the top coverslip contacts the bottom, it traps the molecules in the wells, allowing them to diffuse freely while simultaneously keeping them within the field of view. The sample was kept at 37°C by heaters attached to both the convex and objective lenses. Videos were taken at an exposure of 30 ms once every minute for approximately 1 hour. In between video acquisitions, the top coverslip was raised, allowing new molecules to diffuse into the field of view to replace the previously viewed molecules, and then brought back into contact with the bottom coverslip.

RNPs bound to plasmids were manually counted in each video and then plotted vs time. Bound RNPs were distinguished from free RNPs by differences in their diffusion, size, and intensity. RNPs bound to plasmids appear on video as a slowly diffusing, diffraction-limited spot while free RNPs appear as fast-moving blurs that uniformly illuminate the wells. Clumps of RNPs were ignored in the binding count. Clumps were identified by size and intensity as they were larger and had more fluorophores than the singly labeled bound RNPs. Dark or dim pits outside of the laser beam were excluded from analysis to avoid under-counting molecules. To determine the cut-off position, a Chi-squared test was used to compare the total number of molecules detected in each pit over a dataset (∼60 samplings) to an expected uniformly distributed set of molecules in those same pits. A cut-off radius was selected that optimized the Chi-Squared value while maximizing statistics and excluding any molecules beyond that radius from analysis. Because the pit positions are not constant between datasets, results were normalized per 100 pits.

### Thermal Denaturation Monitored by UV Absorbance

Cas9 alone or assembled as an RNP at 1 µM final concentration (equimolar concentration of all components) was incubated at room temperature for 10 min in degassed 1x UV melt buffer (20 mM Cacodylate, pH 7.5, 150 mM KCl, 1 mM MgCl_2_). Samples were melted in a Cary 400 UV/Vis spectrophotometer at a ramp rate of 1°C/min while UV absorbance at 280 nm was collected every 1 min. Experiments were repeated in duplicate. Melting temperatures were determined using Van’t Hoff calculations and error determined by standard error of the mean using two experimental replicates for each sample.

### Generation of HEK 293T Cells Stably Expressing EGFP

HEK 293T cells stably expressing EGFP (known as EGIP) were a kind gift from Dr. Wen Xue (Yin, H. et al.2017). In addition, we used CRISPR based AAVS1 safe harbor knock-in for generating HEK 293T cells stably expressing an alternative codon-optimized version of EGFP. The EGFP gene was synthesized as a gBlock (Integrated DNA Technologies, IDT) with 15 bp extensions complementary to the ends of pAAVS1-EF1a-BSD-DNR vector (ORIGENE, GE100048) after digestion with KpnI (Thermo Scientific). Insertion was accomplished using the In-Fusion HD Cloning Kit (Clontech/Takara) following the manufacturer’s recommended protocol. Ligated vector was then transformed into NEB stable cells following the manufacturer’s recommended protocol. Single colonies were selected from LB agar-carbenicillin plates and plasmid prepared by midi-prep. A guide sequence targeting AAVS1 locus was synthesized by IDT as two complementary oligonucleotides and annealed to prepare a DNA insert. The pSpCas9 (Addgene 62988) vector was digested with FastDigest BbsI (Thermo Scientific) and the DNA oligo encoding the sgRNA was inserted into the linear vector using T4 DNA ligase (Thermo Scientific) following the manufacturer’s recommended protocol. All clones were verified by Sanger sequencing. HEK 293T cells were co-electroporated (Invitrogen Neon transfection system) with pAAVS1-EF1a-BSD-DNR carrying the EGFP donor template and pSpCas9 expressing sgRNA and Cas9 enzyme. Cells were passaged for three weeks before antibiotic selection to dilute cells containing the donor in non-stable episomal form. Cells were treated with 5 µg/ml of blasticidin for 2-3 days to enrich edited cells. Cells were further selected for EGFP fluorescence with fluorescence assisted cell sorting.

### Cell-based Editing Quantified by Flow Cytometry

HEK 293T cells expressing EGFP versions were grown in Dulbecco’s modified eagle’s medium (DMEM) with 1x non-essential amino acids (NEAA), 5% cosmic calf serum (CCS) and 5% fetal bovine serum (FBS) without antibiotics. HEK 293T cells (3×10^6^) were electroporated (Invitrogen Neon transfection system) with 12 µg (200 ng/5×10^4^ cells) of Cas9 expressing plasmid (LentiV_Cas9_Puro, Addgene, 108100) in a T75 flask and incubated in a 5% CO_2_ incubator for 36 hours in order to allow the plasmid to transiently express Cas9 prior to guide RNA transfection. HEK 293T cells expressing Cas9 and EGFP (5×10^4^ cells) were reverse transfected in four replicates with 10 pmols of crRNA:tracrRNA complex or sgRNA using RNAiMAX (Invitrogen) following the manufacturer’s recommended protocol. Opti-MEM was replaced with full media after 12 hours and cells were allowed to grow for an additional 4 days. Cells were then trypsinized, washed with 1x PBS and resuspended in 200 µl of 1x PBS for flow cytometry analysis (Attune NxT Focusing Cytometer, Thermo Fisher). EGFP was detected using the blue laser (BL1 channel) for at least 20,000 events and analyzed by Attune software (v4.12). A non-transfected control was used for gating (**Extended Data Fig. 20**) and to calculate the total editing efficiency (loss of EGFP) of guide RNAs transfected.

### Cell-based Editing Quantified by rhAmpSeq

Dual guide RNAs were prepared by mixing equimolar amounts of crRNA and tracrRNA in IDT duplex buffer, heating to 95°C and slow cooling to room temperature. Single guide RNAs were prepared by rehydrating in IDT duplex buffer. Ribonucleoprotein complexes were formed by combining the CRISPR-Cas9 nuclease (Alt-R SpCas9 Nuclease V3, IDT) and the guide RNA in a 1:1.2 molar ratio and incubating at room temperature for 30 minutes. RNPs were delivered into Jurkat cells via electroporation as described previously ^33^. Editing events were quantified using the rhAmpSeq system (IDT) as described previously ^33^.

### On Target and Off Target Editing Quantified by Nanopore Amplicon Sequencing

HEK 293T cells expressing EGFP were grown and transfected with Cas9 plasmid and guide RNAs as described above for cell based editing and quantification by flow cytometry. However, 1×10^5^ cells were reverse transfected in three replicates with 40 pmols of crRNA:tracrRNA complex or sgRNA using RNAiMAX (Invitrogen) following the manufacturer’s recommended protocol in a 48-well plate. Opti-MEM was replaced with full media after 12 hours and cells were allowed to grow for an additional 3 days. Cells were then trypsinized, washed with 1x PBS and processed for genomic DNA extraction using Monarch Genomic DNA purification kit (#T3010L) following the manufacturer’s recommended protocol. DNA was quantified using Qubit dsDNA BR kit.

For each sample, PCR reactions were generated with a set of primers designed to generate ∼1,000 bp amplicons with an on or off target Cas9 cut site near the center of the amplicon. PCR was performed in standard Q5 polymerase (New England Biolabs, NEB) conditions, with 0.5 μM final concentration (unless otherwise noted) of primers and approximately 30 ng of genomic DNA/sample. Each PCR reaction was 22 cycles of 98°C for 10 sec, 67°C for 30 sec, and 72°C for 60 sec. Primers for each site are listed below, as well as any deviations from the 0.5 uM final concentration to obtain more equivalent amplicon representation when using multiple primer sets in a single PCR reaction.

Standard nanopore protocol for amplicon barcoding (Native Barcoding Expansion 96 Kit, Oxford Nanopore Technologies, ONT) was followed. Briefly, PCR amplicons are concentrated and purified via SPRI bead cleanup. Amplicons are then ‘end-prepped’ to add a single deoxy adenosine (dA) tail to each fragment with Ultra II enzyme (NEB). Unique barcodes with deoxy thymidine (dT) tails are then ligated to amplicons in each sample. Barcoded amplicons are then pooled and the overhang on the ligated barcodes used to ligate a sequencing adapter. This adapter-ligated and barcoded sample was sequenced on a PromethION P2 solo using MinKNOW (v24.02.16) following standard nanopore protocol. During sequencing and base-calling, quantification of each amplicon in sample is accomplished using RAMPART software ( https://github.com/artic-network/rampart ), with a goal of at least 200x coverage of each amplicon in each sample. After sequencing, percentage of amplicons with Cas9 activity is calculated using the CRISPResso WGS function of the CRISPresso2 software suite ( https://github.com/pinellolab/CRISPResso2 ), then normalized against a control sample with no Cas9 activity. Commands and scripts available upon request.

### *In Vitro* Sequence Preference Determination with Partially Randomized Target DNA and Guide RNA

Partially randomized 200 nt ssDNA ultramers were obtained from Integrated DNA Technologies (IDT). DNA amplification of ssDNA ultramers was conducted using the forward primer pEGIP_5C3_F:/5SpC3/ATGCGATGGAGTTTCCCCA and the reverse primer pEGIP_5C3_R:/5SpC3/CCGCTTTACTTGTACAGCTCG. PCR reactions (25 µL total volume) contained 50 pmol of ssDNA ultramer, 0.5 µM of each primer, 200 µM of dNTPs and 0.02 U/µl of Q5 Hot Start High-Fidelity DNA Polymerase and 1x Q5 reaction buffer (NEB). The PCR cycling conditions were as follows: 98°C for 25 sec, 25 cycles of 94°C for 8 sec, 65°C for 10 sec, and 72°C for 8 sec, followed by a final extension at 72°C for 30 sec. Following PCR amplification, 1 µL (20 U) of Exonuclease I (NEB) was added and incubated at 37°C for 20 min to remove primers and ssDNA. The PCR reactions were bead purified using 3:1 ratio of AMPure XP beads (Beckman Coulter) to reaction volume following the manufacturer’s instructions and eluted in nuclease free water. Purified PCR products were quantified using Qubit 1X dsDNA High Sensitivity (HS) Assay Kit (ThermoFisher Scientific) according to manufacturer’s protocol. The PCR products were visualized on a 2-3% agarose gel stained with ethidium bromide and imaged under UV illumination.

*In vitro* cleavage assays were performed as previously described ^32,43^ with minor changes. Two versions of PCR amplified targets (E2 and E2-Rdm8 or E4 and E4-Rdm8) were used to assess the intrinsic enzyme activity of Cas9 loaded with regular or partially randomized guide RNAs. Briefly, a master mix containing final concentrations of 0.32 µM Cas9, 0.32 µM tracrRNA, 0.32 µM crRNA or 0.32 µM Cas9 and 0.32 µM sgRNA in a 1x cleavage buffer (20 mM Tris-HCl, pH 7.5, 100 mM KCl, 5% glycerol, 1 mM DTT, 0.5 mM EDTA, 2 mM MgCl_2_) supplemented with 0.1 mg/mL of purified yeast tRNA was incubated at 37°C for 10 min to allow the RNP to assemble. Target DNA (2.73 µl of 100 ng/ µL) was added to 77.27 µL of pre-assembled RNP and incubated at 37°C and reactions stopped by phenol-chloroform extraction and ethanol precipitation.

Library preparation was performed using PCR Barcoding Kit (SQK-PCB111.24, ONT) and Native Barcoding Kit 24 V14 (SQK-NBD114-24, ONT). Target DNA cleavage products were end-prepped using 0.75 μL Ultra II End-prep Enzyme mix (NEB), and 1.75 μL Ultra II end-prep reaction buffer (NEB). Reactions were incubated at 20°C for 10 min followed by 65°C for 10 min using a thermal cycler. The PCR products were purified using 3:1 ratio of AMPure XP beads (Beckman Coulter) to reaction volume and eluted in 16 μL of elution buffer (Omega Bio-Tek). Each sample was quantified using Qubit 1x dsDNA High Sensitivity (HS) Kit Assay (ThermoFisher Scientific). For barcode adapter ligation, 15 μL of end-prepped DNA was mixed with 10 μL of the barcode adapter (BCA) and 25 μL of Blunt/TA Ligase Master Mix (NEB) and incubated at room temperature for 30 min. The BCA-ligated DNA was purified using 3:1 ratio of AMPure XP beads (Beckman Coulter) to reaction volume and eluted in 23 μL. The DNA samples were PCR amplified using 1 μL of BC barcode, 1 μL of the forward pEGIP_F primer (0.4 μM) and 25 μL of LongAmp Taq 2X Master Mix (NEB). The PCR cycling conditions were: 95°C for 25 sec, then 13 cycles of 94°C for 8 sec, 62°C for 10 sec, and 65°C for 10 sec, followed by a final extension at 65°C for 30 sec. PCR reactions were purified using 3:1 ratio of AMPure XP beads and eluted in 16 of elution buffer (Omega Bio-Tek). The PCR-amplified samples underwent a second end-preparation as previously described and eluted as previously described in 10 μL of elution buffer (Omega Bio-Tek). The cleaned-up PCR barcoded samples were quantified using Qubit 1x dsDNA High Sensitivity (HS) Assay Kit (ThermoFisher Scientific) and equimolar amount of DNA for each sample was taken forward for native barcoding. Individual end-prepped DNA was barcoded using Native Barcoding Kit 24 V14 (SQK-NBD114-24) by combining 7.5 μL of end-prepped DNA, 2.5 μL of Native Barcode, 10 μL of Blunt/TA Ligase Master Mix (NEB) and incubated at room temp for 20 min. The reactions were stopped by adding 2 μL of 0.5 M EDTA and pooled together. The pooled samples were purified using with 2.8:1 ratio of AMPure XP beads (Beckman Coulter) to reaction volume eluted in 30 μL of elution buffer (Omega Bio-Tek). Adapter ligation was performed by combining 30 μL of pooled barcoded DNA, 5 μL of native adapter (NA), 10 μL of 5x NEBNext Quick Ligation Reaction Buffer (NEB), 5 μL of Quick T4 DNA Ligase (NEB) and incubated at room temperature for 30 minutes. The reactions were purified using 2.2:1 ratio of AMPure XP beads and washed with Short Fragment Buffer (SFB). The final library was loaded on a PromethION R10.4.1 DNA flow cell and sequenced on a PromethION P2 solo using MinKNOW (v24.02.16).

POD5 files were base-called using Dorado (v8.0.1) with the dna_r10.4.1_e8.2_400bps_sup@v5.0.0 model, using the parameters, no-trim and minimum q-score set to 8. The base-called reads were demultiplexed using Dorado demux (v8.0.1) with the SQK-PCB111.24 and no-trim parameter. Custom Python scripts utilizing the Biopython SeqIO module were used to identify reads matching the expected cleaved TCAAGGT and uncleaved TCACCG sequences as well as their reverse complements. For each matched read, 30 nts upstream or downstream of the match were retained as flanking sequence. The reads were subsequently filtered to retain only those of a uniform length and further filtered to retain sequences upstream of the randomized region, containing CACCATGGTGAGGCAA in the forward or TTGCTCACCATGGTG as its reverse complement. Seqtk (v1.4) (https://github.com/lh3/seqtk) was used to generate reverse complements, for consistent orientation in downstream analysis. The filtered forward and reverse DNA sequences were concatenated and then trimmed to retain only the initial 8 nucleotides at the 5’ end of the gRNA binding site on the DNA target. WebLogos of fasta files were generated using WebLogo 2 ^85^ to produce probability-based logos.

### Cryo-EM Structural Studies

Cas9 was assembled with RNA guides and DNA target substrates (ternary complexes) at room temperature for 15 min in 20 mM Tris, pH 7.5, 200 mM KCl, 0.5 mM DTT and 5 mM EDTA. Complexes were prepared at a final of 1.0 to 1.6 mg/mL, placed on ice, and grids prepared within 2-3 h. Cryo-EM grids were Quantifoil 2/2 holey grids, glow discharged prior to vitrification that was carried out on an FEI Vitrobot Mark IV (FEI). Each grid was inspected on a 200-kEv Glacios equipped with a Falcon IV direct electron detector and preliminary data sets were collected for 2-3 hours for each grid with acceptable density of particles and lacking visible aggregates. For both sgE2 and dgE2 complexes, the optimal concentration was found to be at 1.4 mg/ml. Final data sets for the best grids were collected on a 300-kEv Titan Krios G3 cryo-TEM equipped with a GIF BioQuantum 968 energy filter and a Gatan K3 direct electron detector. 5,008 movies were collected for sgE2 complex with pixel size of 0.7 Å, total dose 51.37 e/Å^2^, 49 frames and acquisition time 4.28 s. Two data sets with a total 11,890 movies were collected for the dgE2 complex with similar parameters.

Images were processed with cryoSPARC software. Motion-corrected and ctf-estimated images were curated, resulting in 4,393 movies for the sgE2 complex and 9,251 movies for the dgE2 complex. Several rounds of iterative particle picking using blob and template picking routines followed by the topaz training method and particle selection using 2D classification, *ab initio* modeling, 3D refinement, and 3D classification were performed for each data set. For the sgE2 complex, the best resolution of 3.12 Å was obtained from a final set of 64,788 particles using “Local Resolution” refinement.

In the case of the dgE2 complex, processing of the first data set (5,714 total movies, 4,345 curated movies) yielded a map of 3.31 Å resolution calculated from 105,360 particles using non-uniform refinement. The refined structure represented a “closed” conformation similar to that of the sgE2 complex. Notably, unlike sgE2 data, processing of dgE2 data consistently yielded high contrast 2D classes corresponding to a proposed “open” conformation of the protein alone or in complex with guide RNA only (**Figure S5**). 3D maps for this class were of lower resolution, above 4 Å, which permits placing of major structural domains of Cas9 but not fine nucleic acid detail. In an attempt to improve resolution, we collected a second data set of 6,176 movies from the same grid. 4,906 movies were selected after curation. Additional data provided slightly improve resolution of the map for the closed form to 3.26 Å using 215,418 particles and local refinement. These particles were subjected to 3D classification routine implemented in cryoSPARC v4.1.2. Three identical maps of highest resolution were used as input. One class of 108,048 particles yielded a map of an improved resolution of 3.11 Å. Additional polishing steps of global CTF refinement further improved resolution to 3.03 Å.

Additional data collection did not offer improved resolution of an open form. The highest obtained resolution was of 3.86 Å for 141,467 particles with an overall B-factor of 140 Å^2^.

DeepEMhancer maps were generated to aid modeling and refinement. Models were built and refined using the Refmac program implemented in Phenix combined with manual modeling using Coot. A crystal structure (PDB ID: 4UN3) was used as a starting model for building and refinement of the sgE2 structure. Refined sgE2 structure was modified and used for model building and refinement of the dgE2 structure. PDB ID 7S3H was used for modeling of dgE2 in an open conformation. Only partial RNA strands were modeled into density including nucleotides 40-70 of crRNA in a conformation similar to those of corresponding sgRNA loops in 7S3H, and 7-26 of crRNA and 21-38 of tracrRNA, not previously observed in open conformation complexes. Final structures and experimental data were deposited to PDB and EMBD with PDB ID: 8G1I and EMD-29671 for sgE2, and PDB ID: 8FZT and EMD-29639 for dgE2 in closed conformation, PDB ID: 9B2K and EMD-44111 for dgE2 in open conformation.

### Protein and Nucleic Acid Model Generation

The single and dual RNA-guided Cas9 systems with protein in the catalytically active state were generated for computational studies. The model was prepared in the MODELLER ^86^ using crystal structures in pdb files 5F9R ^53^ and 5Y36 ^54^ as templates. The 5F9R structure corresponds to a precatalytic state of the Cas9, in which the HNH domain is away from the target DNA strand, and in the pseudoactive 5Y36 structure, the HNH domain is very close to the target strand. The precatalytic model of Cas9 cannot capture the structural features associated with the cleavage of the DNA, which is required for the current study. But the nucleic acid strands are more complete in the precatalytic crystal structure. Therefore, a chimeric protein model using the above crystal structures was created and then 5 rounds of steepest descent minimization (5,000 steps) and 5 rounds of conjugate gradient minimization (5,000 steps) were carried out. Five low energy structures generated after minimization were subjected to short 5 ns MD simulations, where harmonic restraints were applied to the protein backbone atoms. Finally, the model with low energy structure corresponding to the catalytic state was used for further studies. The nucleic acid coordinates were obtained from the 5F9R crystal structure. The sequences were changed as per the experimental spacer sequences E1, E2 and E3 in the UCSF Chimera program ^87^. The loop of the single guide system was shortened (by removing 9 base pairs) to match the experimental sequence. To prepare the dual guide system, the first loop was removed (4 bases) and refined from the template nucleic acid structure. The catalytic Mg^2+^ ions were retained from the 5Y36 model. This model generation protocol finally yielded three different models of E1, E2, and E3 sgRNA and dgRNA systems.

The catalytic model of the protein and refined nucleic acid structures were then combined in tleap. The Amber ff14SB ^88^ force field for protein, the parmbsc1 force field ^89^ for DNA, and the ff99bsc0*+χOL3* force field ^90^ for RNA were used. The TIP3P model has been employed for water. The system was solvated using a 10 Å rectangular water box, and NaCl salt concentration of 100 mM was added to the complexes in tleap.

### Molecular Dynamics and Energy Calculations

The molecular dynamics protocol previously reported by Palermo and co-workers ^56^ was used with slight modifications. The system was initially subjected to 10,000 steps (5,000 steps of steepest descent along with 5,000 steps of conjugate gradient) of minimization with a restraint of 300 kcal/mol·Å^2^ on the protein, nucleic acid, and Mg^2+^ ions. Then the restraints were removed, and the complexes were again minimized for 10,000 steps (5,000 steps of steepest descent along with 5,000 steps of conjugate gradient). The time step used for the simulation is 2 fs and SHAKE algorithm was applied to the bonds containing hydrogens. The cut off for long range electrostatic interactions was 10 Å. The heating was done in four stages where systems were heated up from 0 to 50 K and 50 to 100 K by running two NVT ensemble simulations of 50 ps each, imposing restraints of 100 kcal/mol·Å^2^ on the protein, RNA, DNA, and Mg^2+^ ions. The temperature was then increased to 200 K in 100 ps using NVT ensemble in which the restraint was reduced to 25 kcal/mol·Å^2^. Finally, the system temperature was raised to 300 K in an NPT-ensemble simulation of 500 ps without any restraints. 1 ns NPT-ensemble equilibration was done on the system, followed by a test production run of 10 ns NPT ensemble. After observing the 10 ns trajectory of the system, production runs were extended to 100 ns using NVT ensemble in the GPU accelerated version of PMEMD ^91,92^ in AMBER 18 ^93^. The final structure after the 100 ns MD was used as the starting point of the Gaussian Accelerated MD (GAMD) ^94^. A dual boost scheme was used for the GAMD. ∼90 ns of GAMD equilibration followed by 500 ns of GAMD production were carried out for all the six systems. The boost potential was updated every 1.6 ns.

The six trajectories were then analyzed using the CPPTRAJ ^95^ module of AmberTools 19 and VMD 1.9.4. ^96^. The RMSD, RMSF and distance calculations were carried out in the CPPTRAJ module. The cut off for hydrogen bond calculation is 3.5 Å and 135 degrees. The potential of mean force (PMF) is calculated upon accurate reweighting of the simulations using cumulant expansion to the 2^nd^ order. The reweighting was carried out using the PyReweighting toolkit ^97^. The parameters were altered to match our calculations. The PMF calculations were performed on the GAMD trajectories using amino acid distances which were previously determined experimentally and can characterize the catalytic state of the protein. The convergence of the simulation was checked by calculating the PMF plots by varying bin sizes.

The MMGBSA ^98^ calculations with per residue energy decomposition were carried out using the GAMD trajectory, considering every 100 frames. The MMPBSA.py script in AmberTools 19 was used for the purpose. The energy decomposition can be represented as the following:

Energy of Base X = Sum of internal energy of base X + Sum of Interaction with bases + Sum of interaction with amino acid residues (cut-off 6 Å).

Further, difference between the energy of the same base in the single and dual guide system was calculated as:

ΔΔ*E_sg-dg_* = Δ*E*_sg_ − Δ*E*_dg_ ; Δ*E*_sg_ is the per residue energy of the bases in the sg and Δ*E*_dg_ is the per residue energy of the bases in the dg.

The trajectory was visualized in UCSF chimera and the images were rendered using PyMOL 2.4.1.

## ACKNOWLEDGEMENTS

This work was supported by a National Institutes of Health grant (1R01GM135646-01) to K.T.G. and M.J.D and an IRCC IIT Bombay research award grant to P.I.P. We thank Digital Research Alliance Canada for High Performance Computing (HPC) Facilities and Computer Centre IIT Bombay for HPC support. We also thank the Washington University Center for Cellular Imaging for assistance in collecting and analyzing cryo-EM data.

## AUTHOR CONTRIBUTIONS

R.C. performed all biochemical and cell-based experiments, analyzed results, prepared figures, and assisted with manuscript writing. S.S. performed all MD simulations, analyzed MD results, prepared figures, and assisted with manuscript writing. A.A.P. designed and performed guide randomization and nanopore sequencing experiments, analyzed results, prepared figures, and assisted with manuscript writing. M.S.B. designed primer sets and optimized nanopore-based sequencing analysis of editing at genomic on- and off-target sites, analyzed results, prepared figures, and assisted with manuscript writing. F.S. performed CLiC experiments, analyzed results, prepared figures, and assisted with manuscript writing. A.K. synthesized and purified chemically modified tracrRNAs. S.H. assisted with GaMD simulations, initial energy plots and comparison circles, and helped write the manuscript. R.T. performed editing experiments at endogenous loci using modified and unmodified guide RNAs from Integrated DNA Technologies, analyzed results, and assisted in preparing figures. M.J.D. supervised nucleic acid synthesis and assisted with manuscript preparation. S.L. supervised CLiC experiments and assisted with manuscript preparation. S.K. analyzed all cryo-EM data, prepared figures, and assisted with manuscript preparation. P.I.P. supervised MD simulations, analyzed MD data, prepared figures, and assisted with manuscript preparation. K.T.G. conceived and organized the project, supervised all biochemical and cell-based experiments, interpreted results, prepared figures, and wrote the manuscript.

## INCLUSION and ETHICS

Research was conducted in an actively inclusive manner to incorporate diverse ideas and perspectives amongst researchers and within the research environment to ensure equitability and avoid potential biases. Experiments, analyses, and interpretation of results were conducted with high regard for academic integrity and reproducibility during the research process.

## DATA AVAILABILITY

All data supporting the findings of this study are available within the paper and its Extended Data. Raw sequencing data from nanopore analyses and MD simulation files are available upon request or available in Zenodo with accession number XXX. Cryo-EM structures have been deposited in the RCSB Protein Databank with accession numbers 8G1I, 8FZT, and 9B2K.

---

**figure S1.**
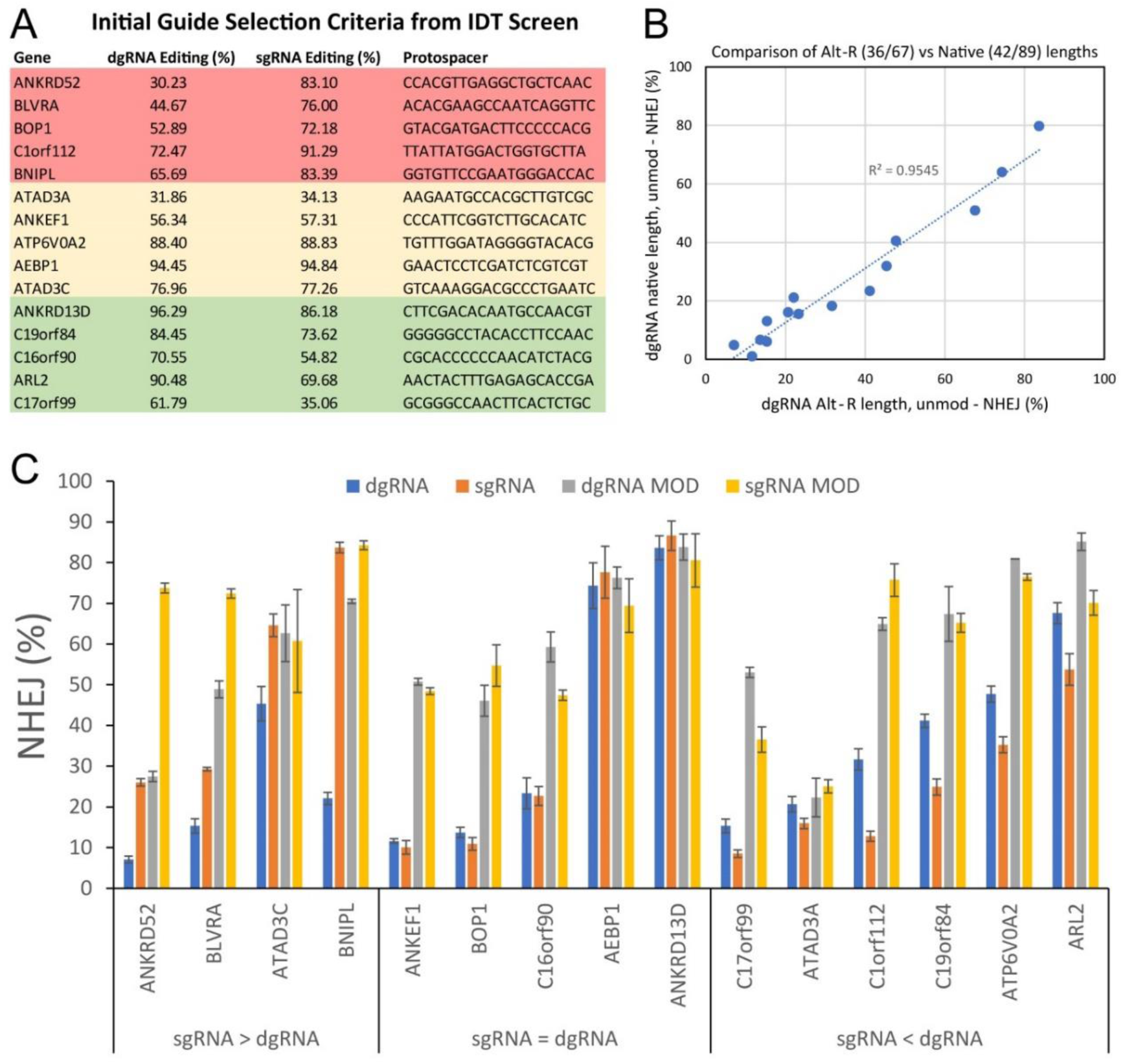
Editing of select endogenous genes with native and chemically modified sgRNA- and dgRNA-guided Cas9. (**A**) Fifteen guides targeting endogenous genes were initially selected based on performance in a screen using a commercially available AltR-CRISPR system. Five guides were selected where sgRNAs outperformed dgRNAs, five where dgRNAs outperformed sgRNAs, and five where sgRNA and dgRNA performed equally well. (**B**) sgRNAs and dgRNAs were resynthesized as native, unmodified guides and the native unmodified dgRNAs compared to chemically modified AltR-CRISPR dgRNAs. Chemical modification (AltR-CRISPR), proprietarily designed to increase stability of the crRNA-tracrRNA repeat and resist nuclease degradation, generally improved the gene editing performance of all 15 guides with high correlation (R^2^ = 0.9545). (**C**) Comparison of gene editing activity for all 15 guides as native and chemically modified (MOD) (AltR-CRISPR) sgRNAs and dgRNAs. Guides are grouped based on performance of native (unmodified) sgRNA versus native (unmodified) dgRNA. Error is reported as standard deviation (3 experimental replicates).

**figure S2.**
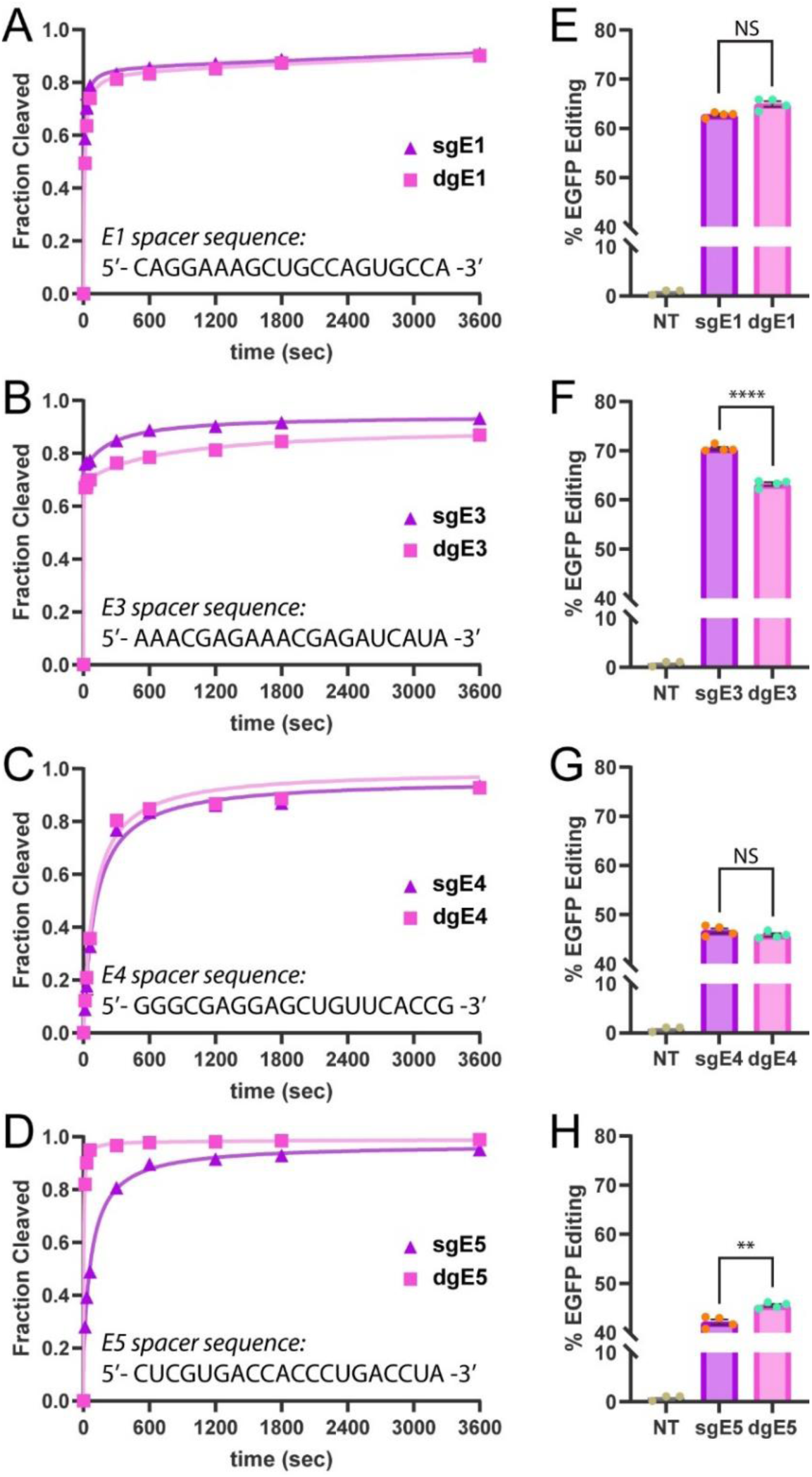
Catalytic activity of Cas9 guided by sgRNA and dgRNA bearing E1, E3, E4, and E5 spacer sequences. (**A**-**D**) *In vitro* time course cleavage assays for E1, E3, E4, and E5 spacer sequences (2 replicates). (**E**-**H**) Cell-based EGFP editing assays for E1, E3, E4, and E5 spacer sequences (4 replicates). Unpaired t-test p-values, ** = 0.0016, **** = <0.0001, NS = not significant.

**figure S3.**
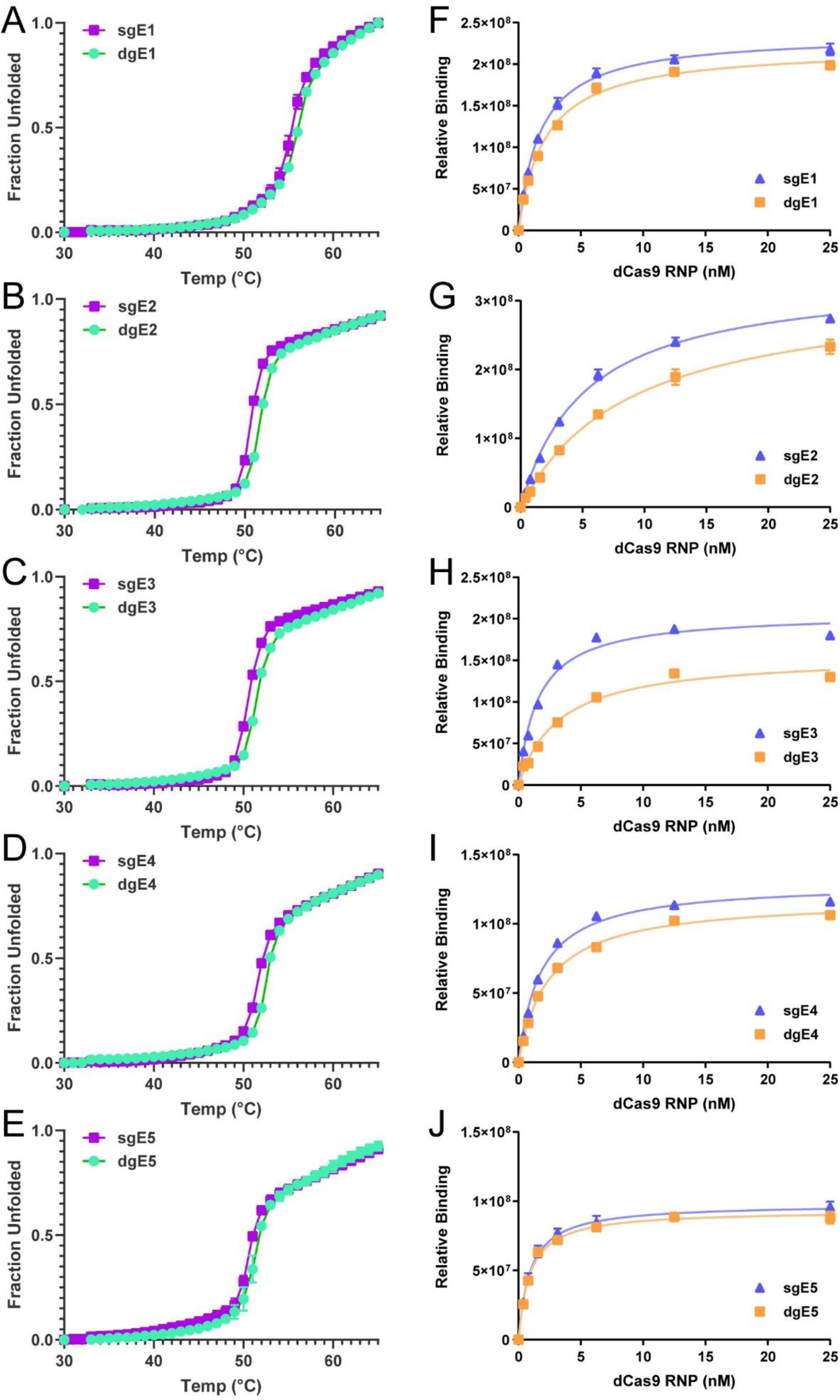
Thermal stability and target DNA binding of Cas9 RNPs guided by sgRNA and dgRNA bearing E1, E3, E4, and E5 spacer sequences. (**A**-**D**) RNP thermal stability measured by absorbance at 280 nm (2 replicates). (**E**-**H**) dsDNA target binding affinity measured by dot blot equilibrium binding (2 replicates).

**figure S4.**
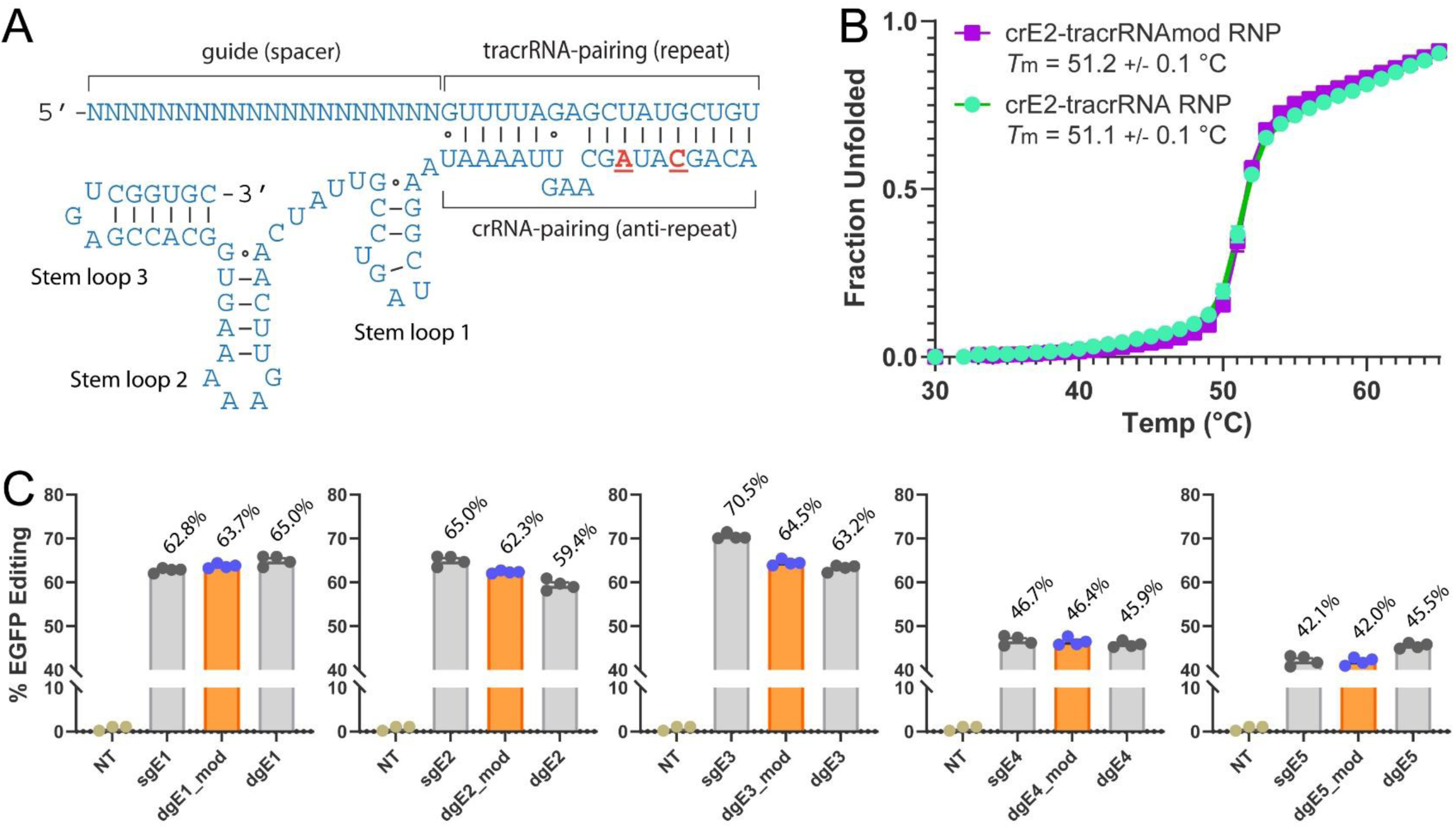
Thermal stability and gene editing activity of dgRNA-guided Cas9 with locked nucleic acid modifications in the tracrRNA repeat sequence. (**A**) Schematic representation of dgRNA. Locked nucleic acid (LNA) modifications are indicated by red, bold, underlined nucleotides. (**B**) Thermal stability of Cas9 RNPs assembled with E2 crRNA and modified and unmodified tracrRNA measured by absorbance at 280 nm (n=2). Range or values reported at +/−. (**C**) Cell-based EGFP editing assays for dgRNAs with chemically modified tracrRNA (tracrRNAmod) (n=4). Gene editing percentages shown above bar graphs are reported as the mean. Error is S.E.M.

**figure S5.**
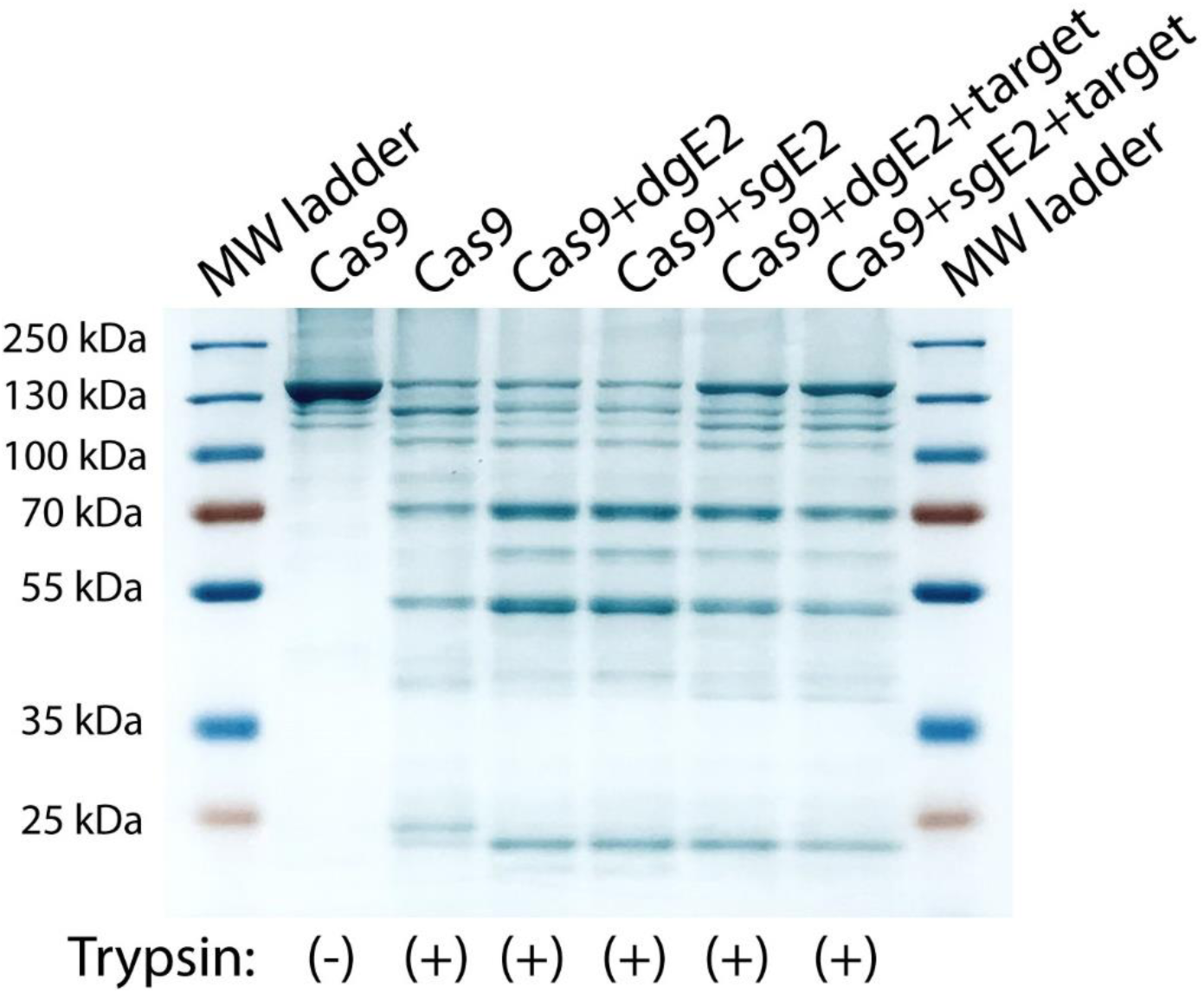
No major differences are observed in trypsin hydrolysis cleavage patterns between Cas9s assembled with dgE2 and sgE2 or bound to E2 DNA target. (**F**) Resolution of limited trypsin hydrolysis products by SDS-PAGE for Cas9 in RNP and ternary (target bound) complexes.

**figure S6.**
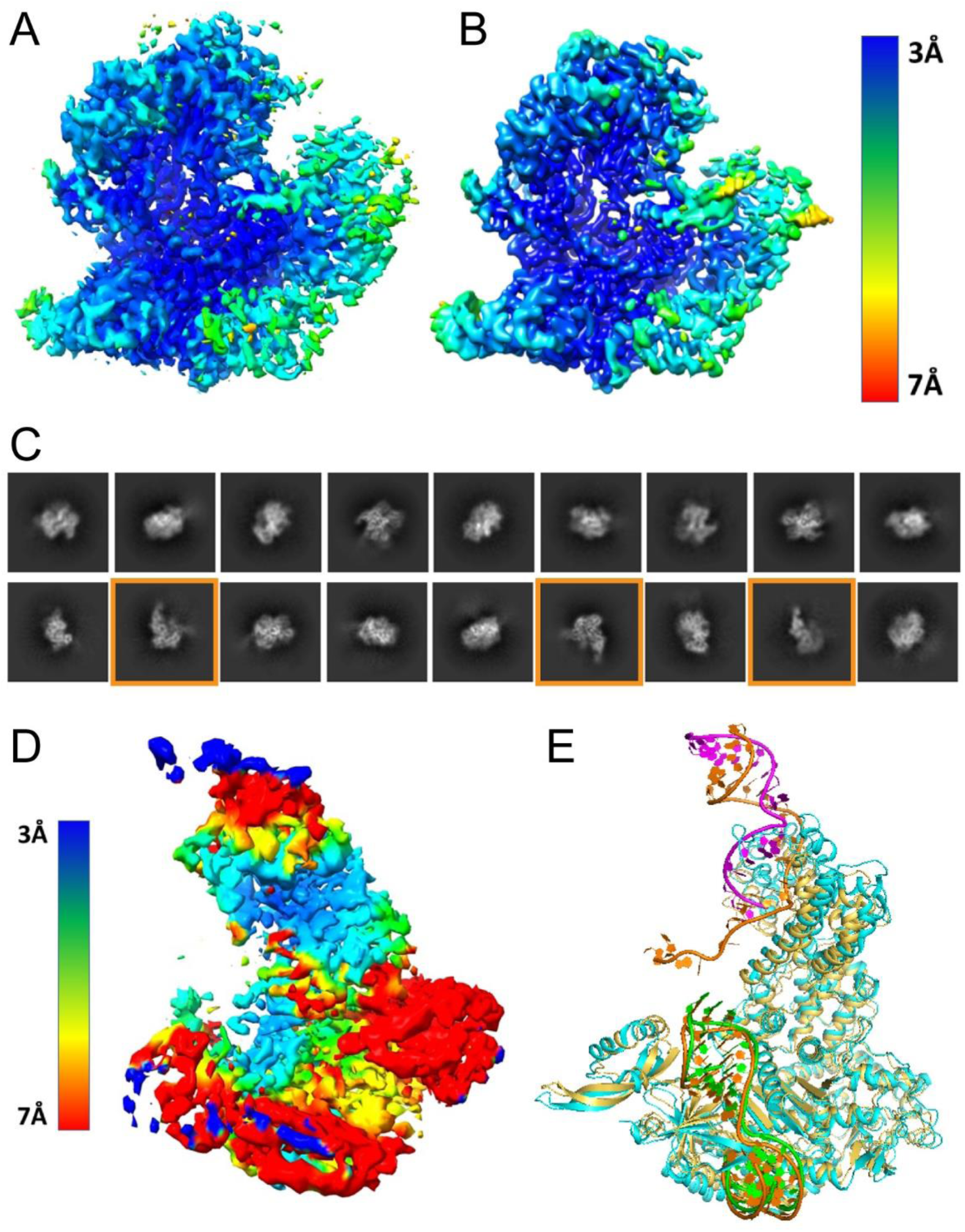
Electron density resolution and 2D classification of single and dual guide Cas9 ternary complexes solved by cryo-EM. Final electron density maps for closed forms of sgE2 (**A**) and dgE2 (**B**) ternary complexes solved by cryo-EM 3D reconstruction color-coded by local resolution of 3-7 angstroms. (**C**) 2D classifications for dgE2 Cas9 ternary complexes. 2D classifications enclosed in orange boxes are classified as “open” complexes. (**D**) low resolution 3D reconstruction of a dgE2 Cas9 ternary open complex color-coded by local resolution of 3-7 angstroms. (**E**) Fitting of a Cas9 “open” structure previously solved by Doudna and colleagues (PDB ID: 4CMP) to the low resolution structure reconstructed in panel D (PDB ID: 9B2K).

**figure S7.**
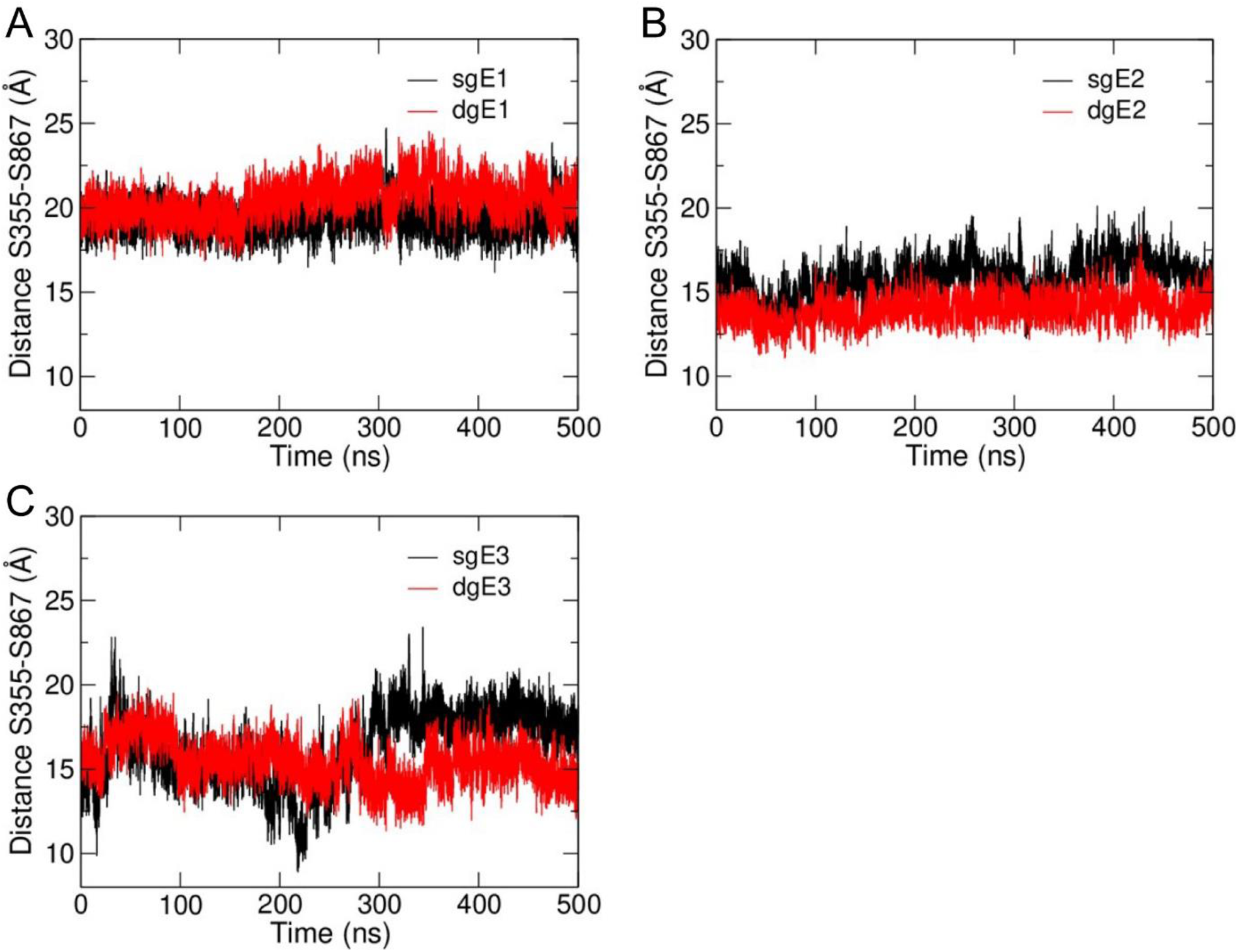
Time dependent distance plots of the S355-S867 amino acids in modeled sgRNA and dgRNA Cas9 ternary complexes with E1 (**A**), E2 (**B**), and E3 (**C**) spacer sequences from the 500 ns GAMD trajectories.

**figure S8.**
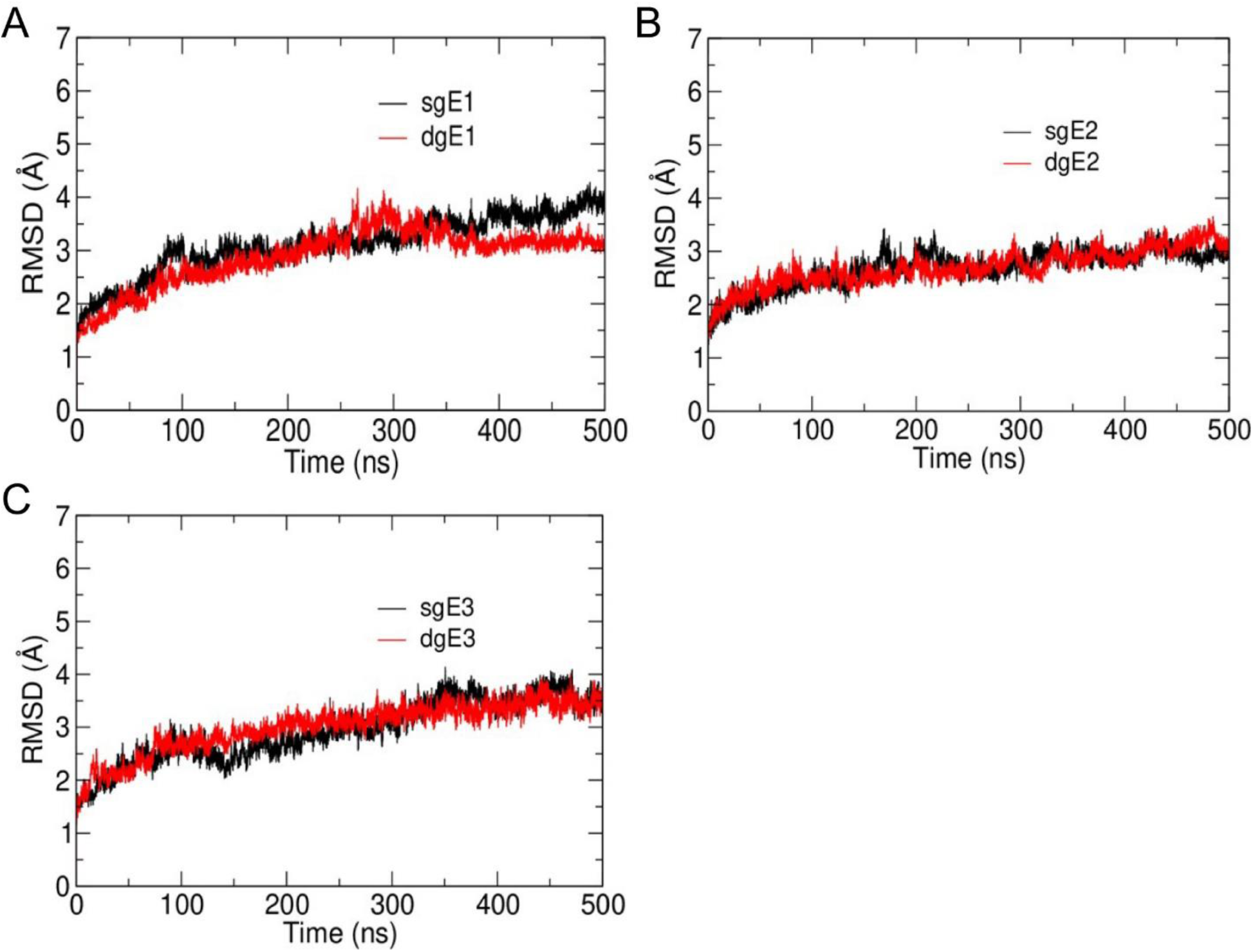
RMSD graphs of the protein backbone in modeled sgRNA and dgRNA Cas9 ternary complexes with E1 (**A**), E2 (**B**), and E3 (**C**) spacer sequences. The RMSD values were calculated from the 500 ns GAMD trajectories.

**figure S9.**
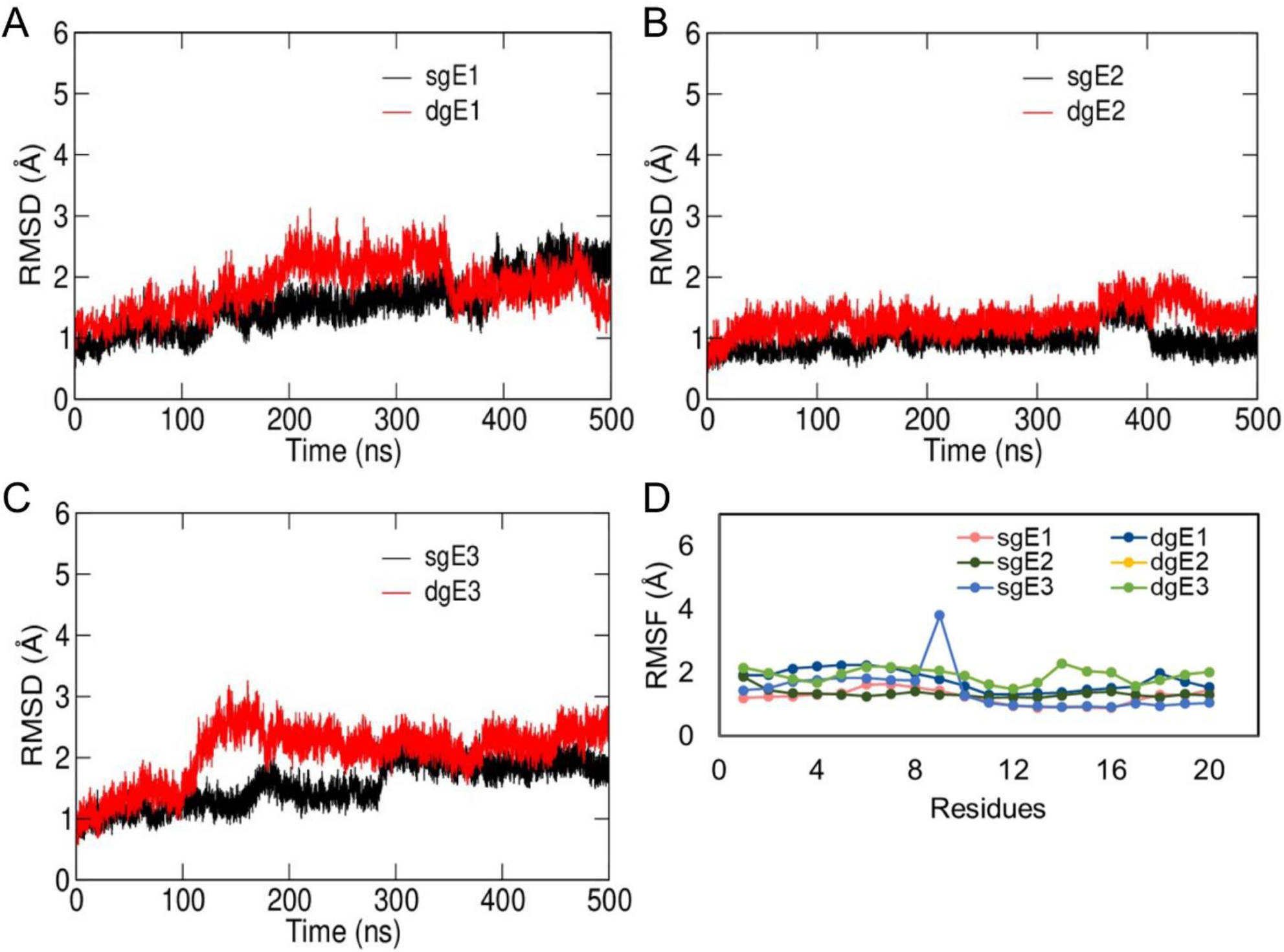
RMSD graphs of the guide RNA backbone in modeled sgRNA and dgRNA Cas9 ternary complexes with E1 (**A**), E2 (**B**), and E3 (**C**) spacer sequences. The RMSD values were calculated from the 500 ns GAMD trajectories. Only the first 20 nucleotides were considered for the plots shown. (**D**) Per nucleotide RMSF of the guide RNA in modeled sgRNA and dgRNA Cas9 ternary complexes with E1, E2, and E3 spacer sequences

**figure S10.**
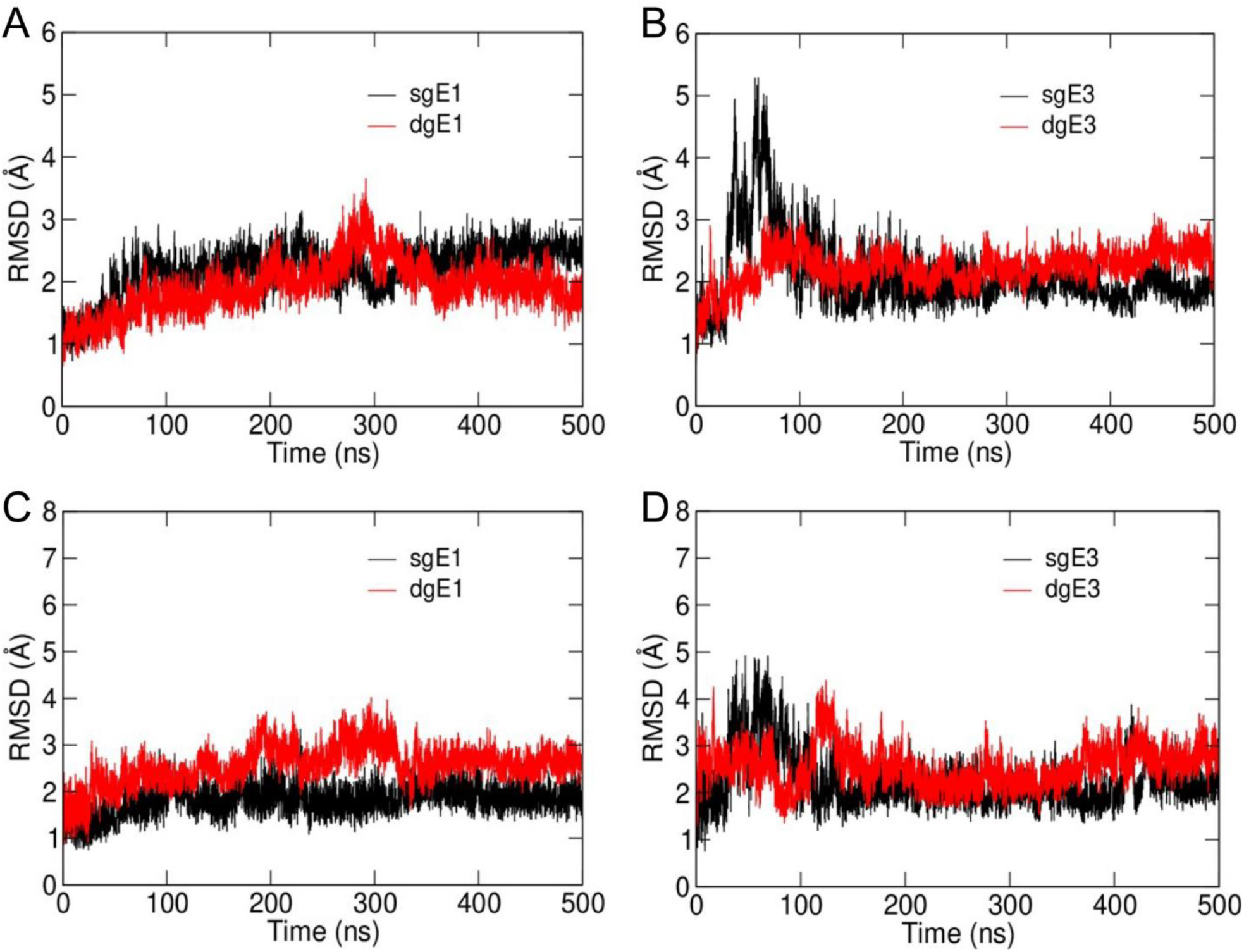
RMSD graphs of the DNA backbone in modeled sgRNA and dgRNA Cas9 ternary complexes for the target strand (TS) of E1 (**A**) and E3 (**B**) spacer sequences and for the non-target strand (NTS) of E1 (**C**) and E3 (**D**) spacer sequences. The RMSD values were calculated from the 500 ns GAMD trajectories.

**figure S11.**
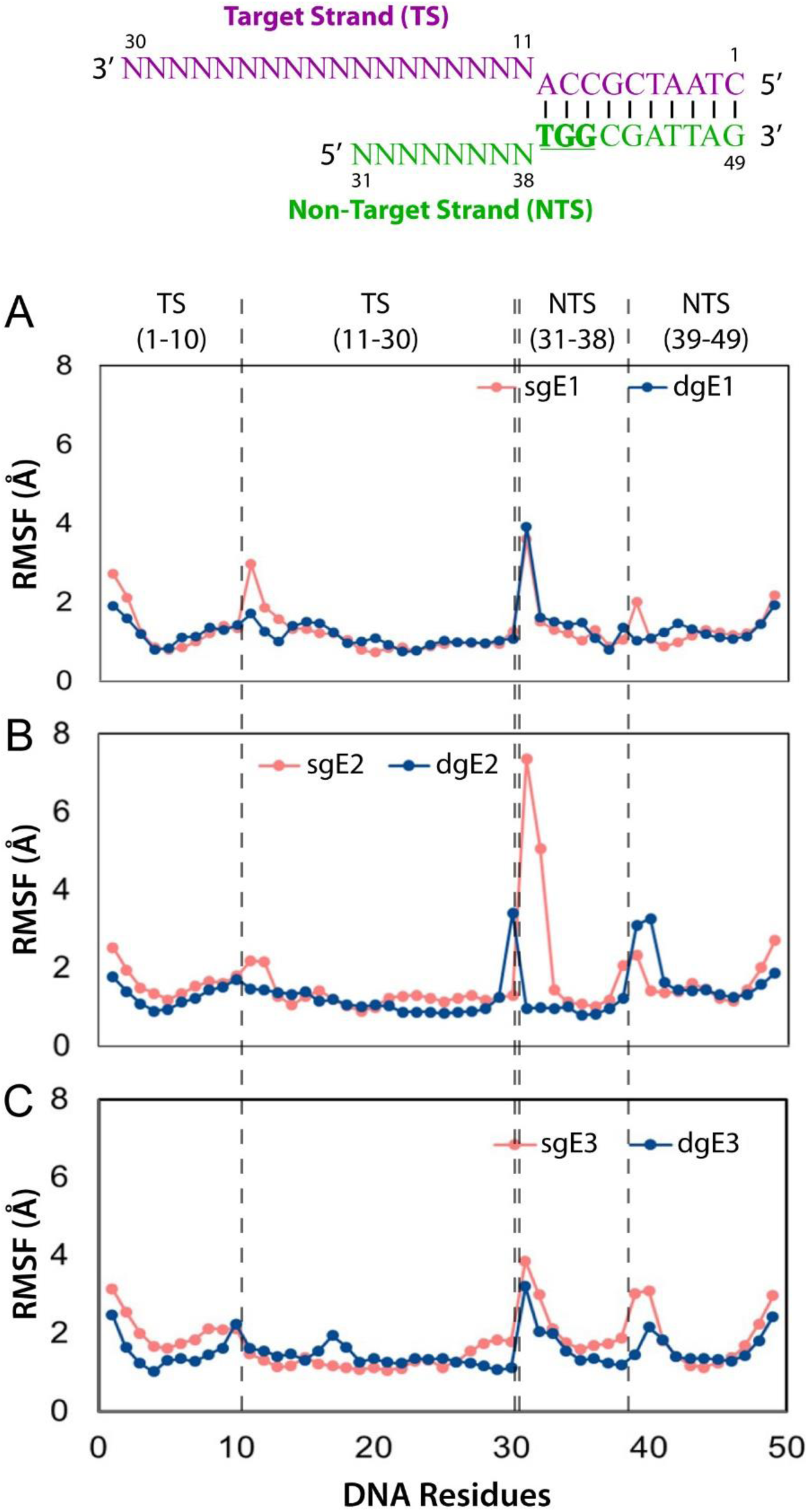
Per nucleotide RMSF of the TS and NTS DNA in modeled sgRNA and dgRNA Cas9 ternary complexes with E1 (**A**), E2 (**B**), and E3 (**C**) spacer sequences.

**figure S12.**
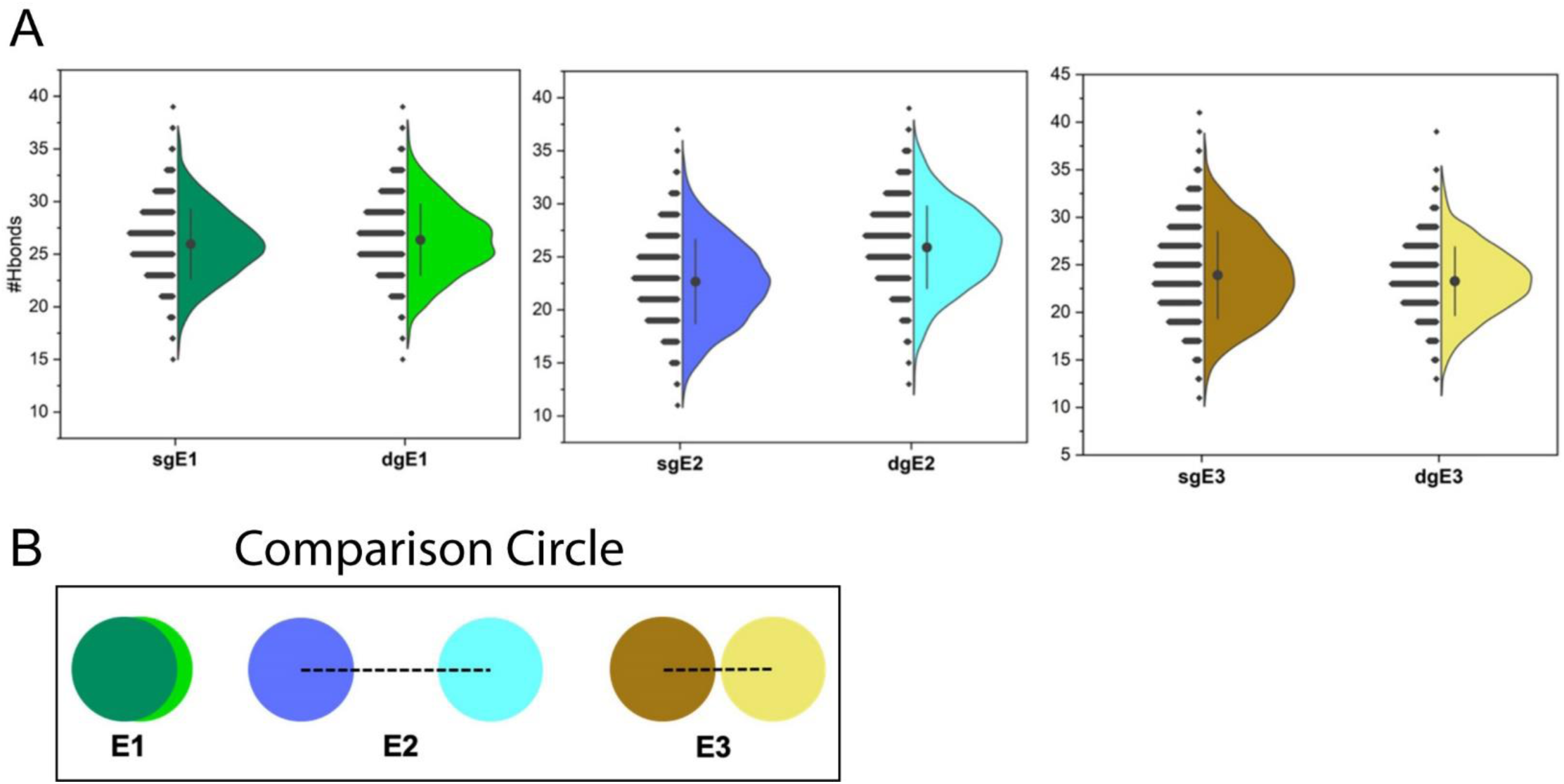
(**A**) Number of Hydrogen bonds between target DNA and Cas9 were calculated in modeled sgRNA and dgRNA Cas9 ternary complexes with E1, E2, and E3 spacer sequences. The values were calculated in VMD using every 100 frames from the 500 ns trajectory. The central point represents the mean value of the histogram distribution. (**B**) The comparison circle was generated using the various parameters from the MD simulation trajectories. The image shows the deviation between E1, E2, and E3 sgRNA and dgRNA cas9 ternary complexes. sgE1, dgE1, sgE2, dgE2, sgE3, and dgE3 are represented in dark green, light green, purple, cyan, brown, and yellow, respectively. The plot is obtained by using the comparison circle algorithm with the RMSD of the DNA strands, interdomain angles, and H-bond counts as the input.

**figure S13.**
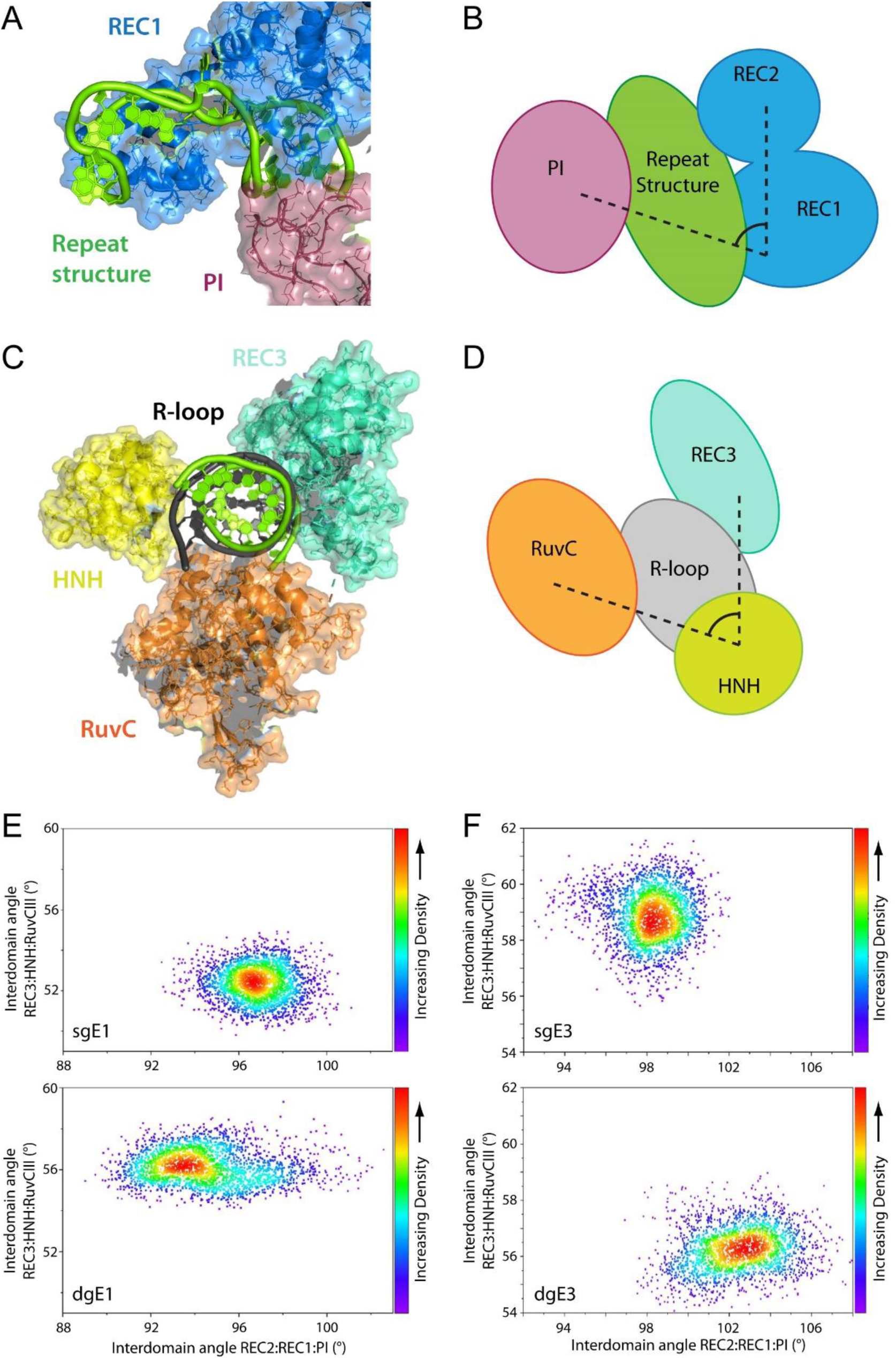
Structural rendering of sgE2 Cas9 cryo-EM illustrating the close contacts of (**A**) the REC1 and PI domains with the repeat structure and (**B**) the REC3, RuvC, and HNH domains with the R-loop structure. (**C-D**) Illustrations of the interdomain angle measurements used for preparing density plots. (**E**) Density plots between the REC3:HNH:RuvCIII and REC2:REC1:PI domains of sgE1 and dgE1 as well as sgE3 and dgE3 ternary complexes.

**figure S14.**
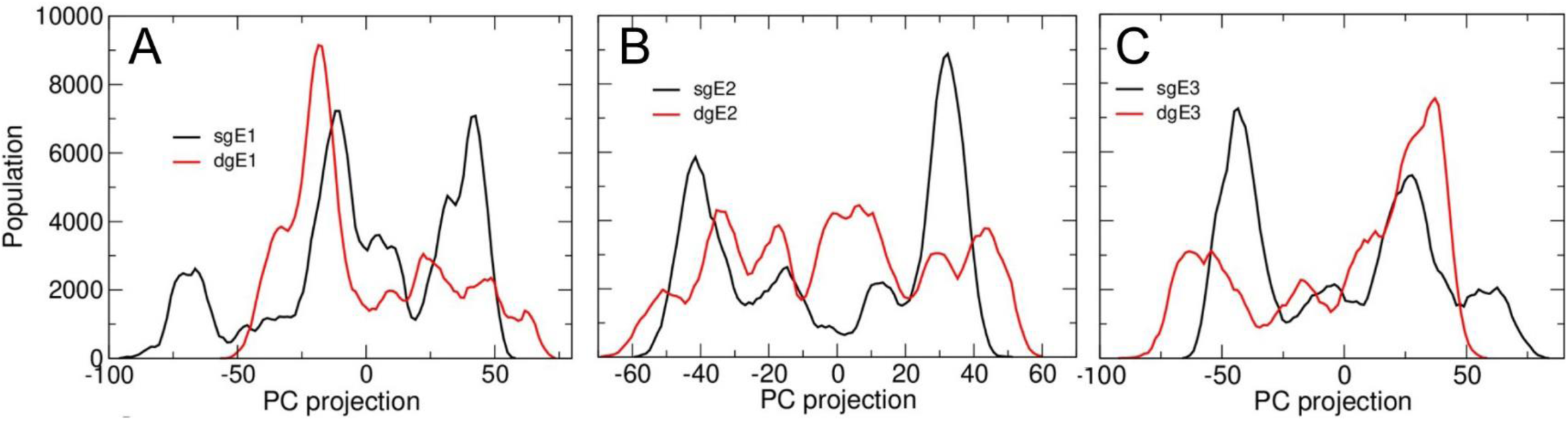
Histogram distribution showing the projection of PC1 of the protein backbone of (**A**) E1, (**B**) E2, and (**C**) E3 Cas9 sgRNA and dgRNA ternary complexes. All calculations have been performed using the 500 ns GaMD trajectories.

**figure S15.**
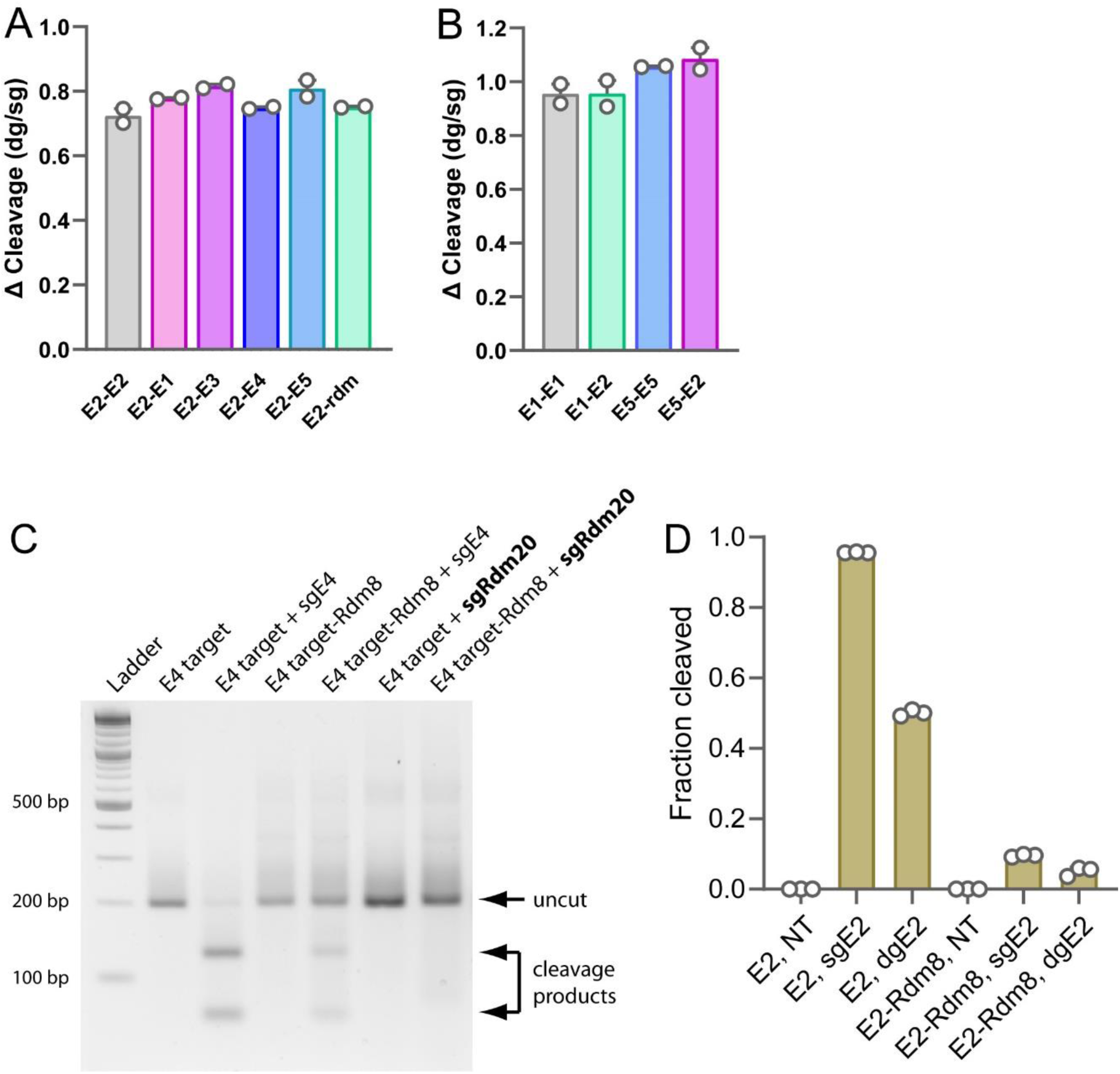
(**A-B**) Quantification of data shown in Figure 4A-B demonstrating the average difference in *in vitro* cleavage by Cas9 guided by E2, E1, or E5 guides with target DNA substrates containing a matched TS but unmatched and non-complementary NTS sequences (n=2). (**C**) Separation of *in vitro* cleavage products on an agarose gel. Target DNA is either E4 or E4 with the first 8 nts of spacer target region randomized (E4 target-Rdm8). Guides used were either perfectly complementary sgRNA (sgE4) or sgRNA with all 20 spacer nts randomized (sgRdm20). (**D**) Quantification of *in vitro* cleavage products from agarose gel separation. Target DNA is either E2 or E2 with the first 8 nuts of the spacer target region randomized (E2-Rdm8). Guides used were either perfectly complementary sgRNA (sgE2) or dgRNA (dgE2) (n=3

**figure S16.**
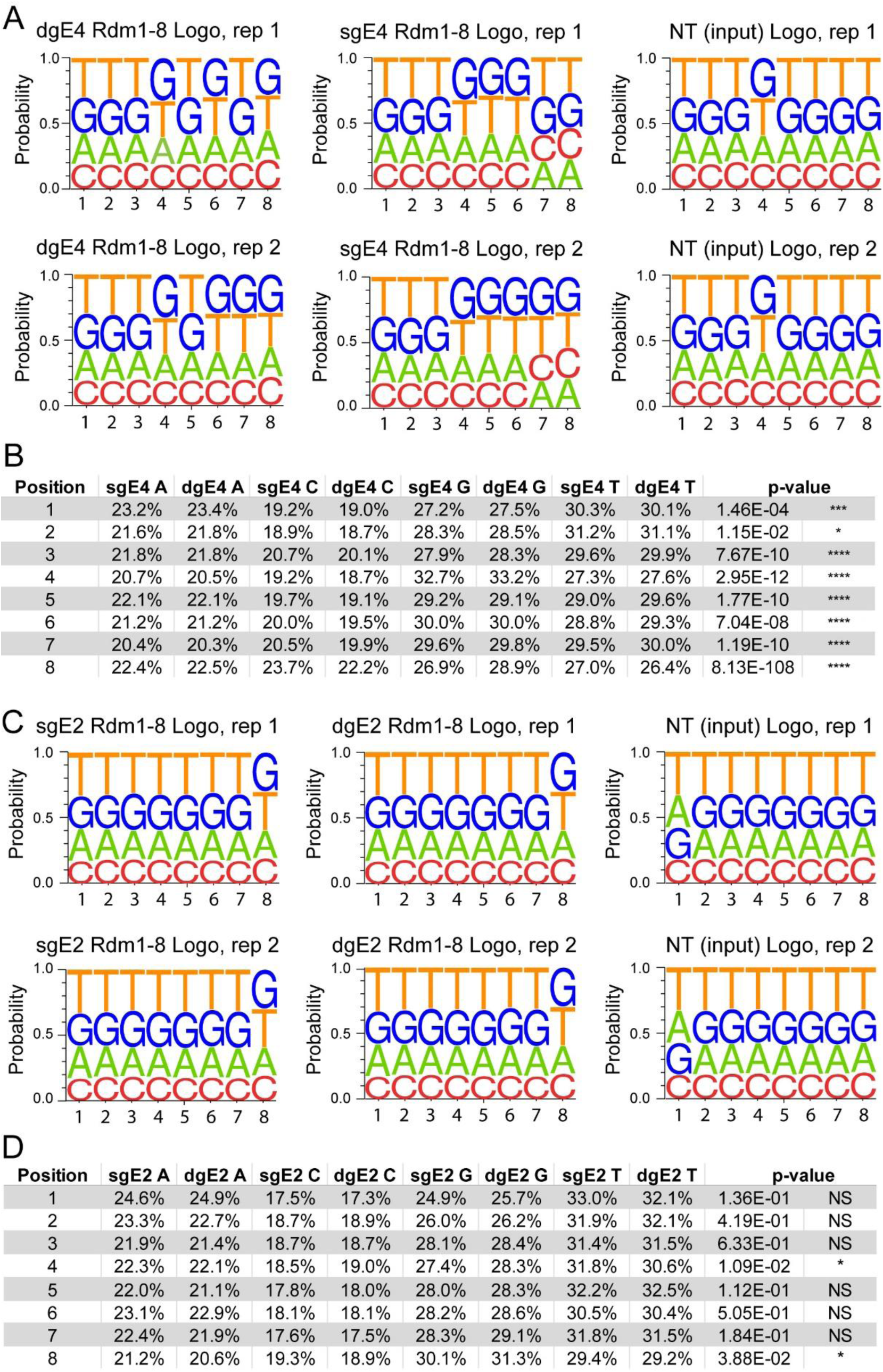
Experiments to determine sequence preferences induced by sgRNAs versus dgRNAs using nanopore sequencing of *in vitro* cleavage products. (**A**) Sequence logos generated for replicates 1 and 2 after nanopore sequencing of E4-Rdm8 target cut with partially randomized dgE4 and sgE4. (**B**) Breakdown of average percent occurrence of individual nts at each position from nanopore sequencing and statistical significance of results at each position. (**C**) Sequence logos generated for replicates 1 and 2 after nanopore sequencing of E2-Rdm8 target cut with partially randomized dgE2 and sgE2. (**D**) Breakdown of average percent occurrence of individual nts at each position from nanopore sequencing and statistical significance of results at each position.

**figure S17.**
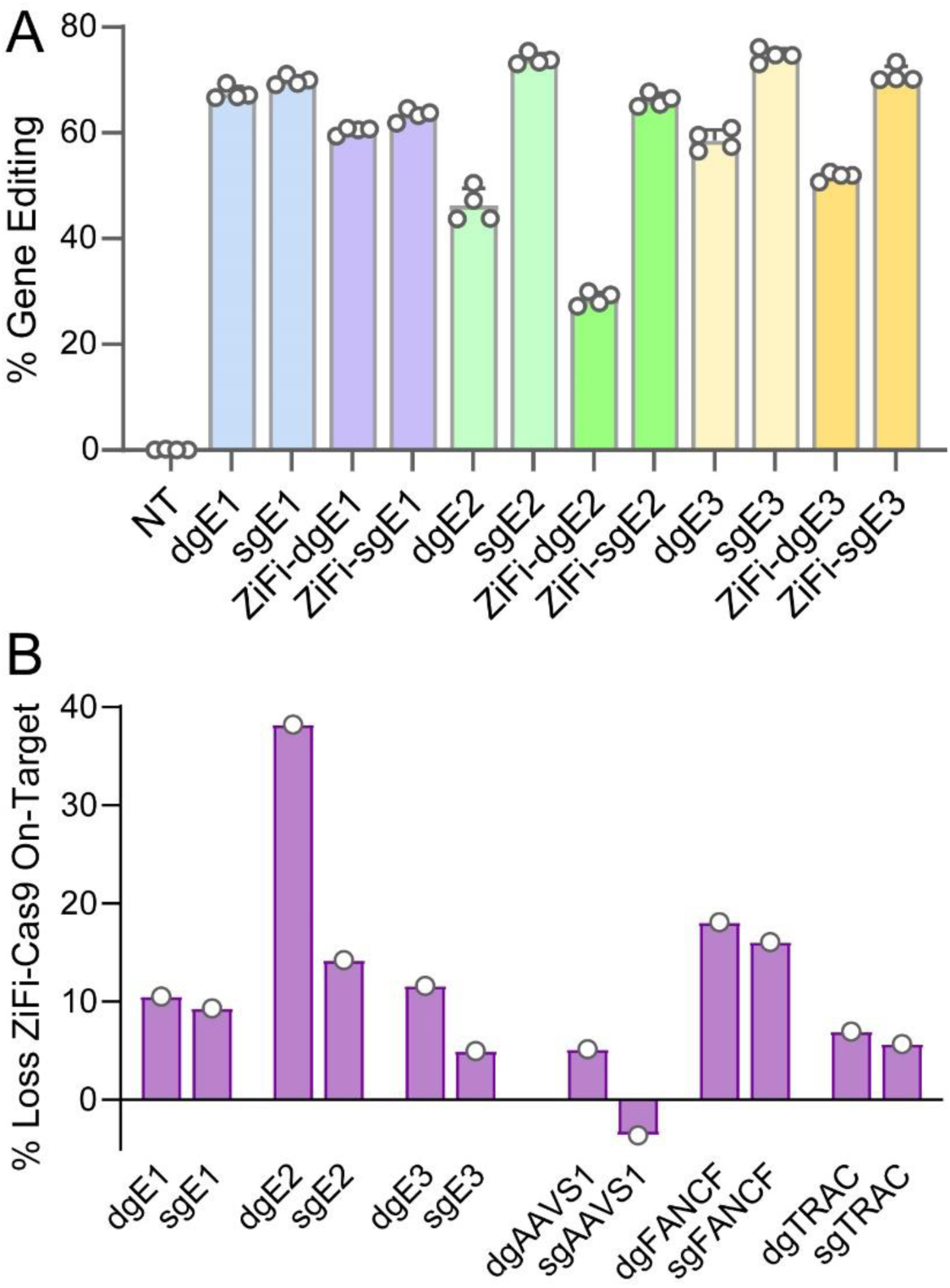
On-target and off-target editing with ZiFi-Cas9 and repeat-truncated sgRNA. (**A**) Editing by E1, E2, and E3 dgRNAs or sgRNAs with normal or ZiFi-Cas9 quantified by EGFP knockout in HEK 293T cells and flow cytometry. Error is S.E.M. (**B**) Loss of average on-target editing for results from panel A and results shown in Figure 4E.

**figure S18.**
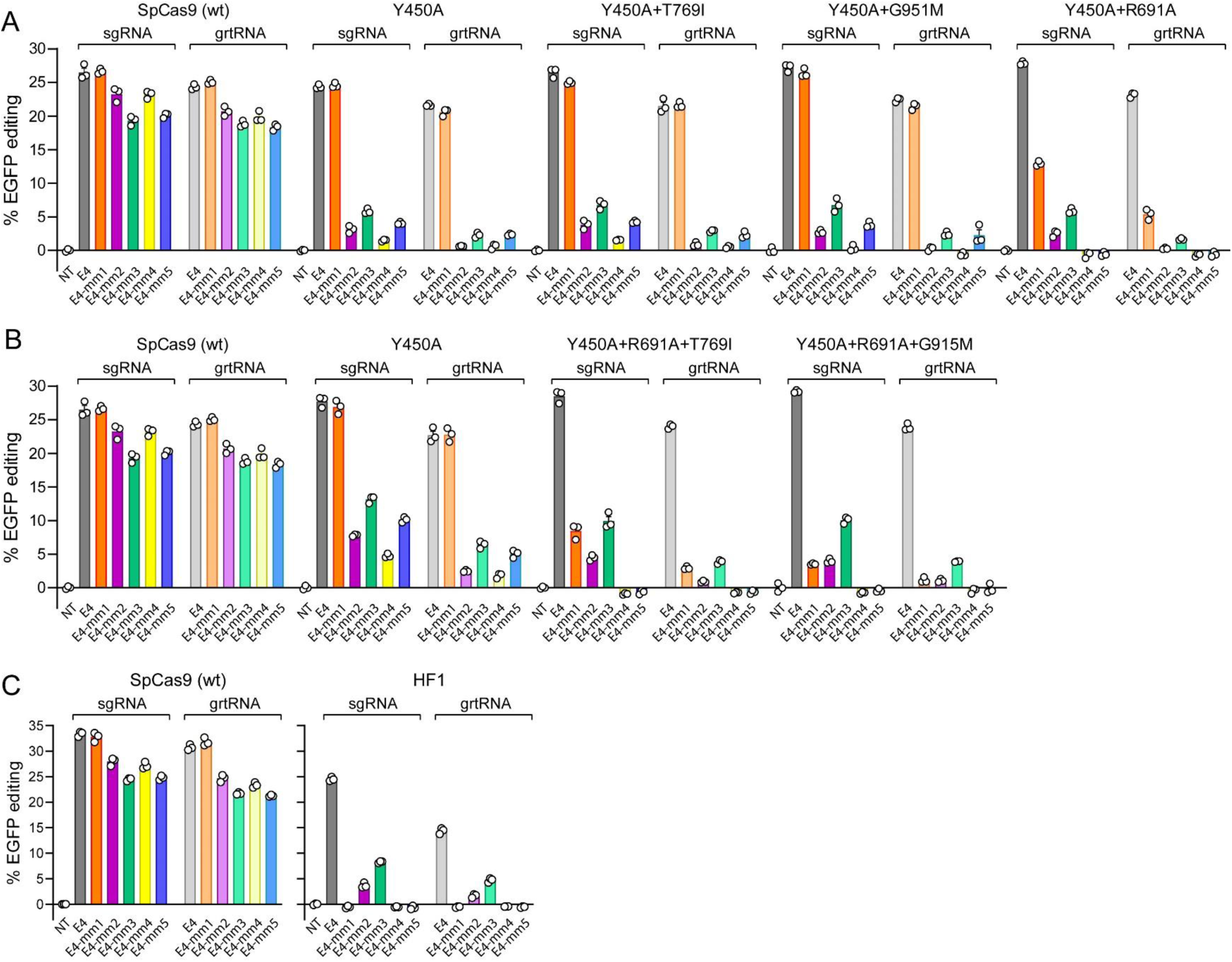
On and off target editing of EGFP with E4 guides and high-fidelity mutants of Cas9. (**A-B**) Screening of specific ZiFi mutations to identify which ones contribute the most to ZiFi-Y450A performance using EGFP editing and E4 guides (n=3). Error bars are S.E.M. (**C**) Testing of HF1 on and off target editing using EGFP editing and E4 guides (n=3). Error bars are S.E.M.

**figure S19.**
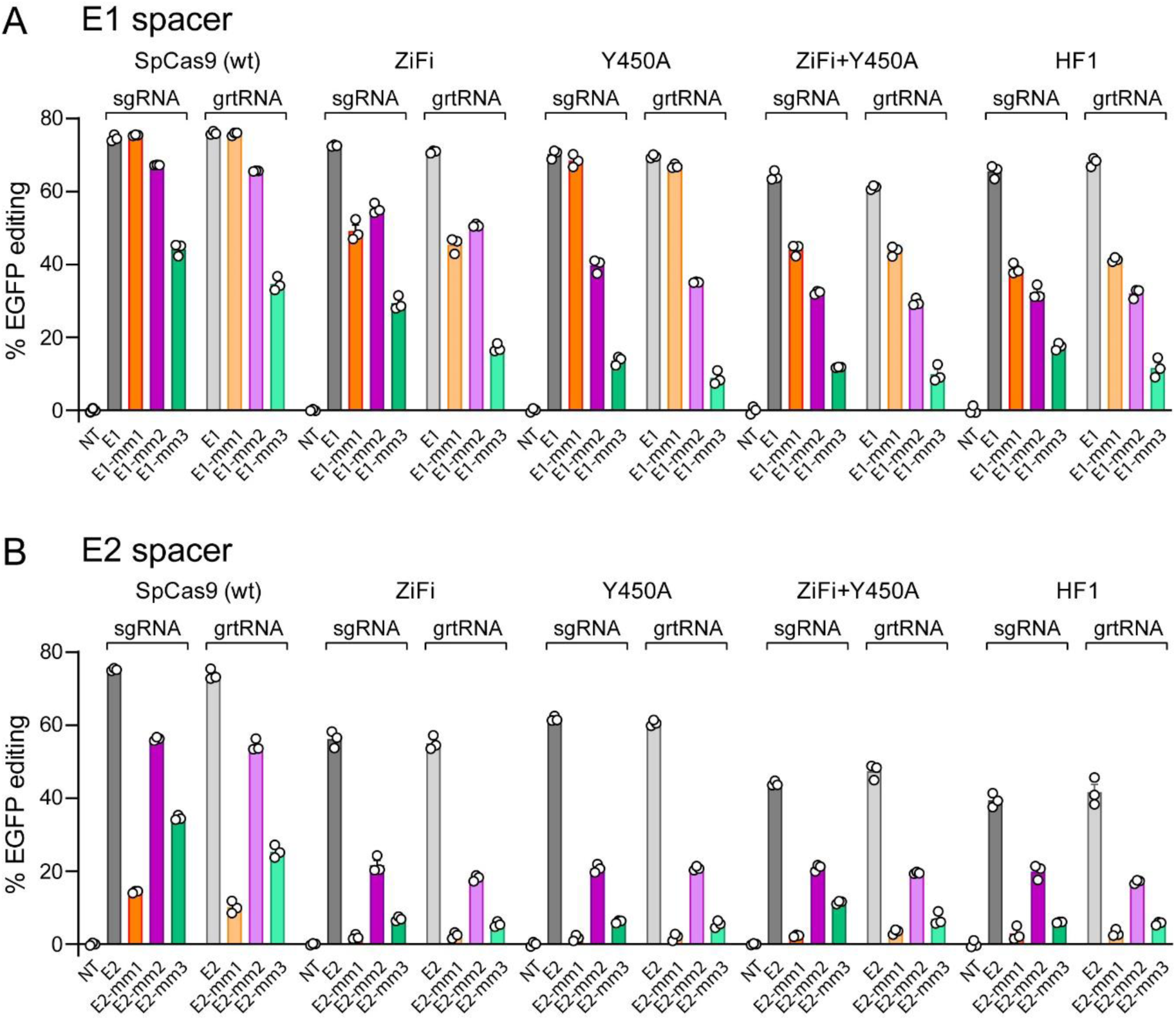
On and off target editing of EGFP with E1 and E2 guides and high-fidelity mutants of Cas9. (**A-B**) Editing of EGFP with the E1 and E2 spacer and mismatch-containing guides and high-fidelity Cas9 variants measured by flow cytometry (n=3). Error bars are S.E.M.

**figure S20.**
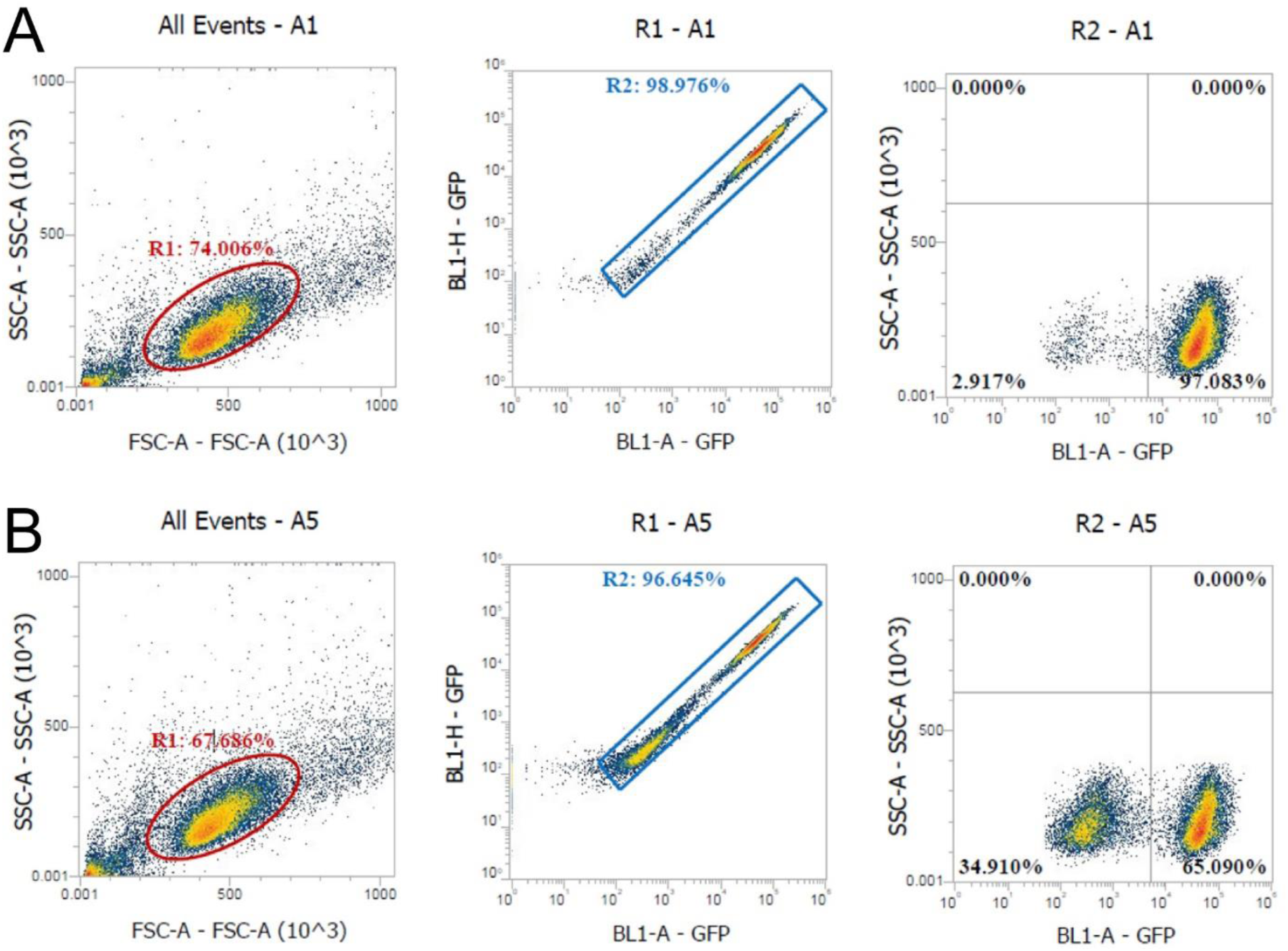
Representative gating and example data for flow cytometry quantification of EGFP editing in HEK 293T cells. Flow cytometry of (**A**) non-treated and (**B**) guide RNA transfected HEK293T cells. Gate 1: forward vs. side scatter (FSC vs SSC) density plot (for identifying cell population of interest and excluding debris). Gate 2: BL1-Area vs. BL-1-Height density plot (for excluding doublets). Gate 3: BL1-A vs. SSC-A two parameter density plot (for distinguishing EGFP positive cells and non fluorescent cells).

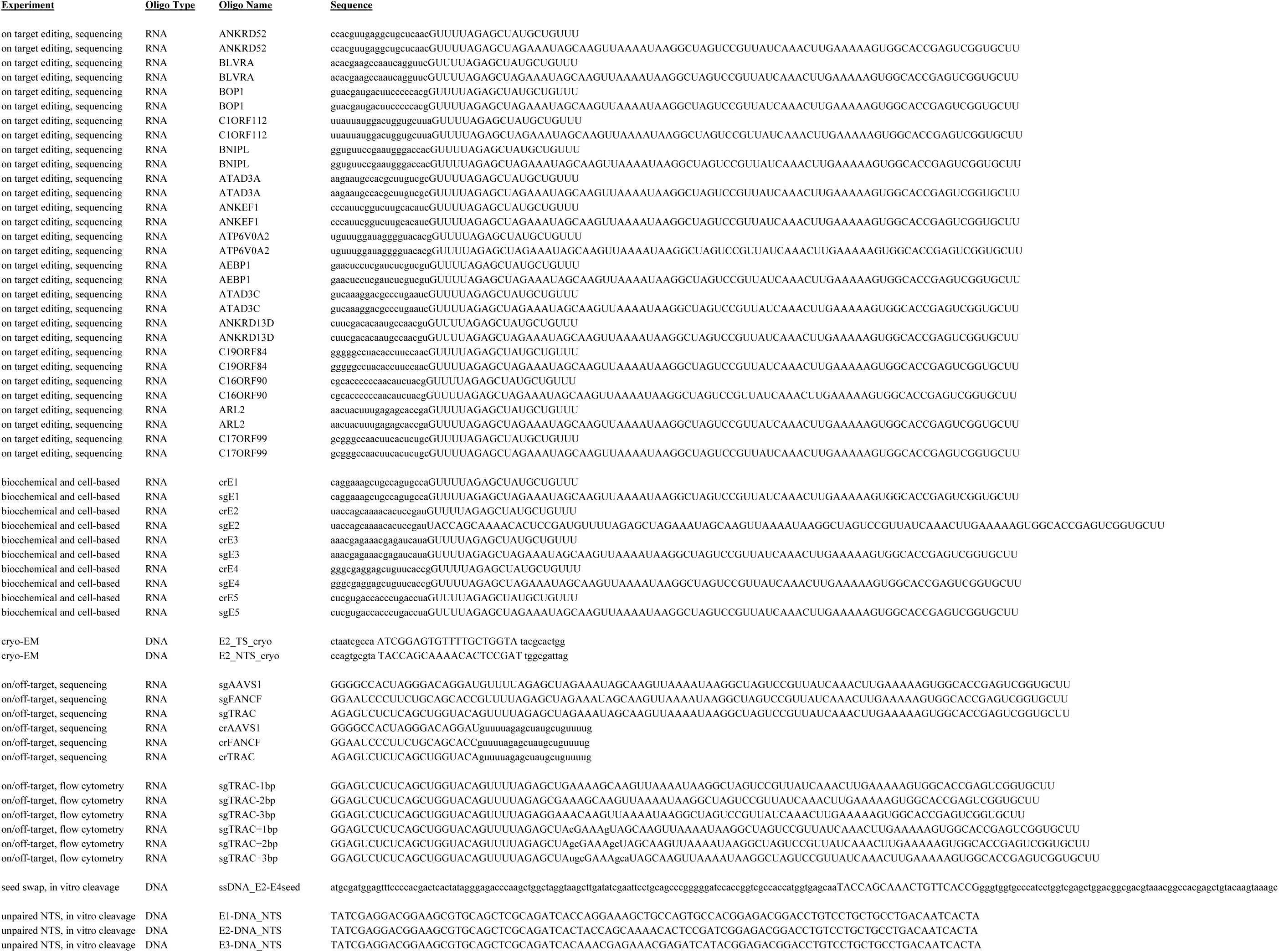
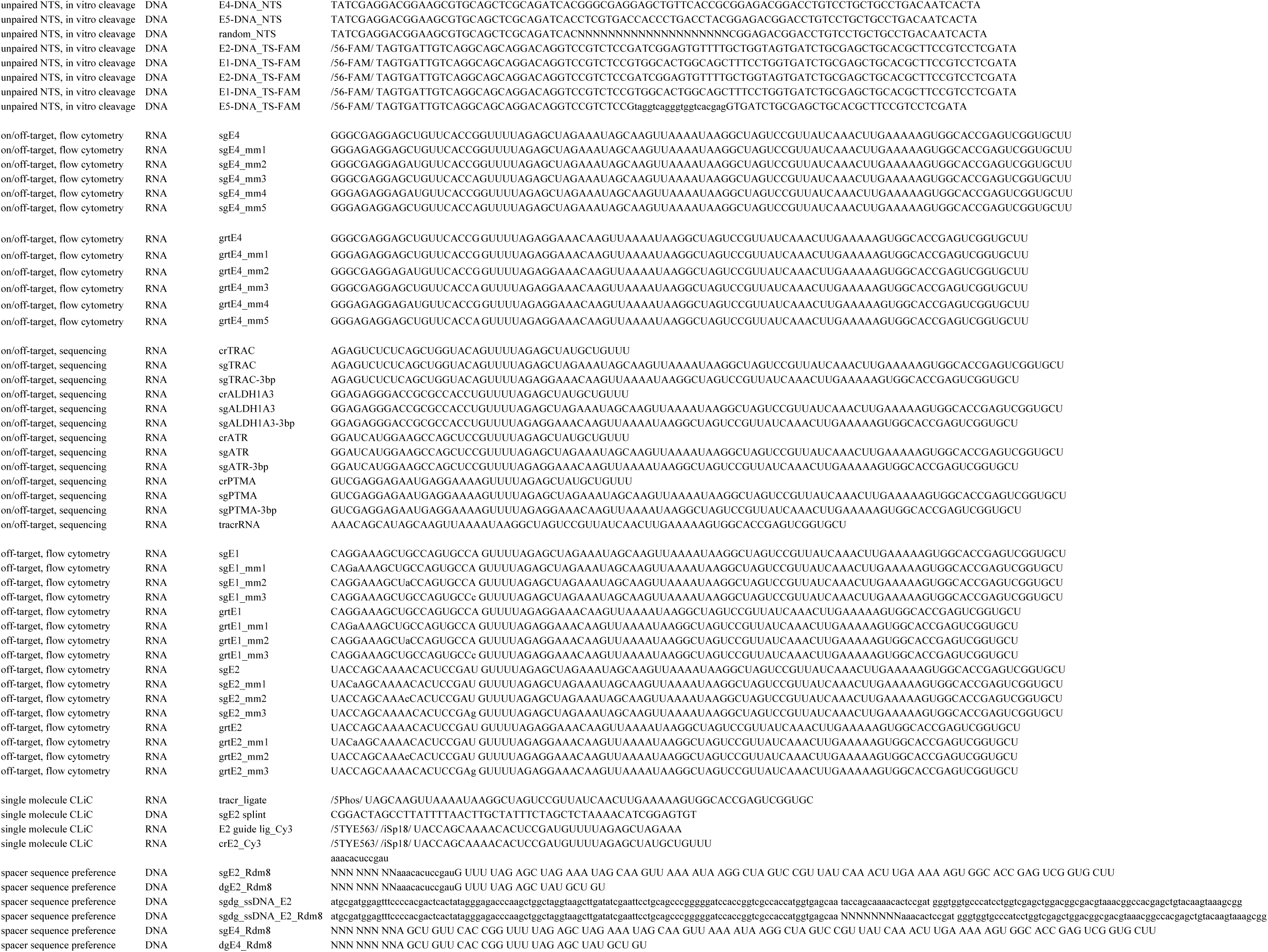
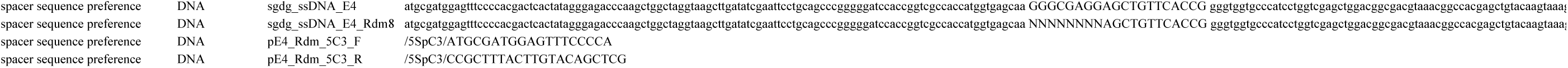

